# A circuit mechanism for the coordinated actions of opposing neuropeptide and neurotransmitter signals

**DOI:** 10.1101/2022.08.03.502700

**Authors:** Marta E. Soden, Joshua X. Yee, Larry S. Zweifel

## Abstract

Fast-acting neurotransmitters and slow, modulatory neuropeptides are commonly co-released from neurons in the central nervous system (CNS), albeit from distinct synaptic vesicles^1^. The mechanisms of how co-released neurotransmitters and neuropeptides that have opposing actions, e.g., stimulatory versus inhibitory, work together to exert control of neural circuit output remain unclear. This question has been difficult to resolve due to the inability to selectively isolate these signaling pathways in a cell- and circuit-specific manner. To overcome these barriers, we developed a genetic-based anatomical disconnect procedure that utilizes distinct DNA recombinases to independently facilitate conditional *in vivo* CRISPR/Cas9 mutagenesis^2^ of neurotransmitter- and neuropeptide-related genes in distinct cell types in two different brain regions simultaneously. With this approach we demonstrate that the stimulatory neuropeptide neurotensin (Nts) and the inhibitory neurotransmitter *γ*-aminobutyric acid (GABA), which are co-released from neurons in the lateral hypothalamus (LH), work coordinately to activate dopamine neurons of the ventral tegmental area (VTA-DA). We show that GABA release from LH-Nts neurons acts on GABA neurons within the VTA to rapidly disinhibit VTA-DA neurons, while Nts signals through the Nts receptor 1 (Ntsr1) on VTA-DA neurons to promote a slow depolarization of these cells. Thus, these two signals act on distinct time scales through different cell types to enhance mesolimbic dopamine neuron activation, which optimizes behavioral reinforcement. These data demonstrate a circuit-based mechanism for the coordinated action of a neurotransmitter and neuropeptide with opposing effects on cell physiology.

---

Numerous cell types within the CNS co-release combinations of neuropeptides and neurotransmitters with opposing actions on cellular physiology^3-5^. A major unresolved question is how these opposing signals work to coordinate circuit function. An example of this paradoxical co-release is neurons in the LH that release stimulatory Nts and inhibitory GABA. The majority of Nts-producing neurons in the LH, like most LH afferents to the VTA, are reported to be GABAergic^6-8^, and LH-Nts neurons are proposed to activate dopamine-producing neurons within the VTA to regulate behavioral reinforcement^8-15^, but it remains unclear what the roles of Nts and GABA are in this process. We propose a circuit mechanism whereby GABA release from LH-Nts neurons acts on local VTA-GABA neurons to rapidly disinhibit VTA-DA cells, while Nts acts through Ntsr1 on VTA-DA neurons to provide a slow stimulatory depolarization.

Consistent with previous reports^6,7^, we found that the majority of Nts neurons in the LH are GABAergic, as evidenced by co-expression of *Nts* and *Slc32a1* (*Vgat*, the vesicular GABA transporter) (Fig. 1a-c, Supplementary Table 1). To establish the connectivity of LH-Nts neurons in the VTA, we generated double transgenic mouse lines that express Cre-recombinase from the Nts locus (*Nts*-Cre) and Flp-recombinase from the dopamine-selective tyrosine hydroxylase locus (*Th*-Flp; Extended Data Fig. 1a,b) or the Vgat locus (*Vgat-*Flp). We injected *Nts-*Cre::*Th-* Flp or *Nts-*Cre::*Vgat-*Flp mice with a Cre-conditional adeno-associated virus (AAV) containing channelrhodopsin 2^16^ (AAV1-FLEX-ChR2-eYFP) into the LH and a Flp-conditional AAV encoding the fluorescent protein mCherry (AAV1-FLEXfrt-mCherry) into the VTA to mark cells for slice electrophysiology (Figure 1d). Light activation of LH-Nts inputs to the VTA generated outward inhibitory currents in VTA-GABA neurons, but we found minimal inhibitory connectivity with VTA-DA neurons (Fig. 1e-f). We detected small inward excitatory currents in fewer than 5% of VTA-GABA or VTA-DA neurons, consistent with the low expression of *Slc17a6* (*Vglut2*) in LH-Nts neurons (Fig. 1a-f).

**Figure 1.**
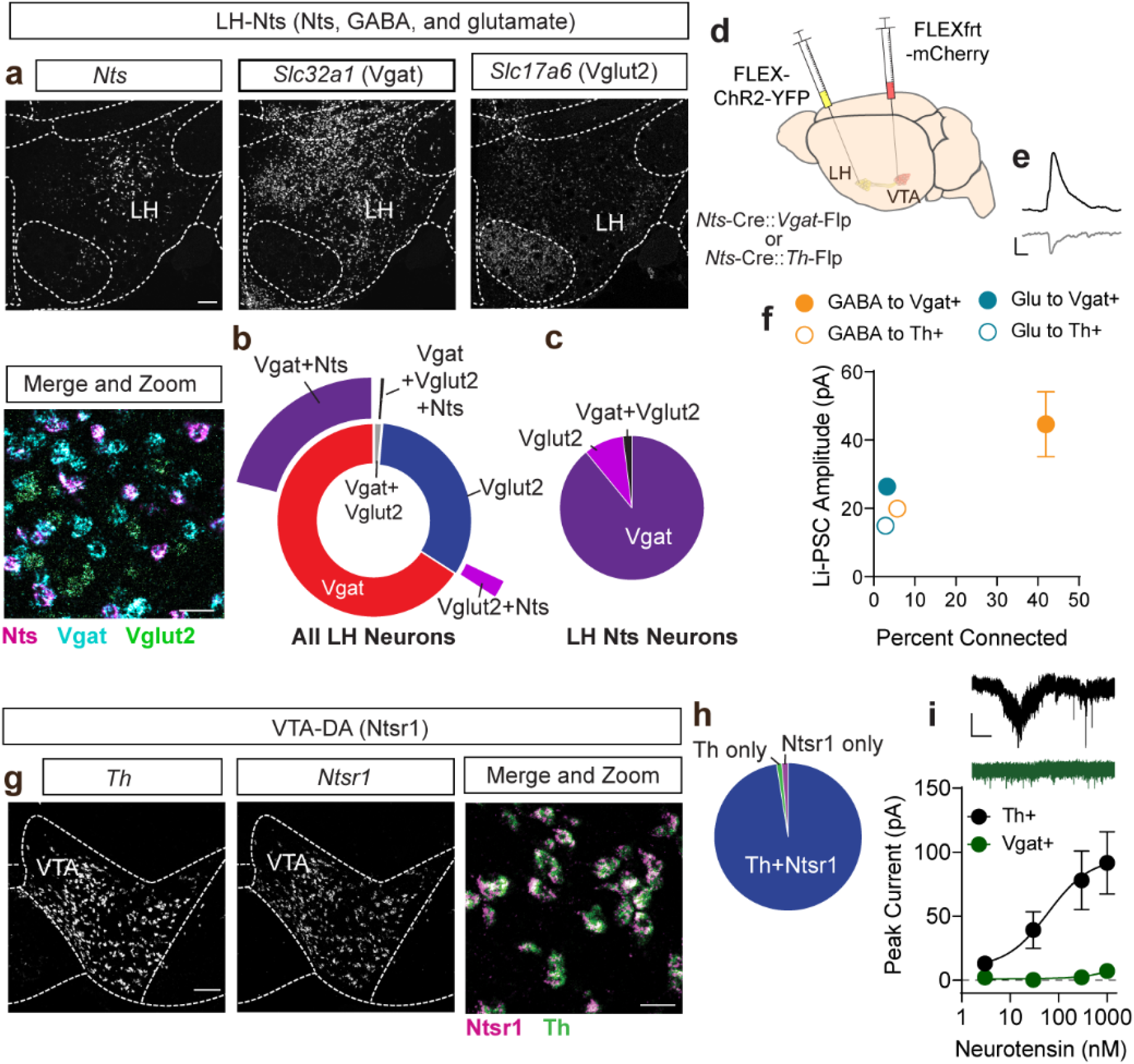
Connectivity of LH-Nts neurons and Ntsr1 signaling in the VTA. (**a**) *In situ* hybridization in the LH for *Nts, Slc32a1* (Vgat), and *Slc17a6* (Vglut2). Scale bars = 150 µm (greyscale images) and 25 µm (zoom). (**b**) Quantification of all Nts, Vgat, and Vglut2 expressing neurons in the LH. (**c**) Quantification of Vgat and Vglut2 expressing Nts neurons of the LH. (**d**) Schematic of viral injection strategy for slice electrophysiology. (**e**) Example traces of inhibitory (top) and excitatory (bottom) light-evoked postsynaptic currents (Li-PSCs) onto VTA neurons (scale bar = 25 pA, 10 ms). (**f**) Percent of recorded neurons with a detectable Li-PSC and average Li-PSC amplitude recorded from VTA Vgat+ or Th+ neurons (GABA to Vgat n=13/31 cells, GABA to Th n=2/35 cells, Glu to Vgat n=1/31 cells, Glu to Th n=1/35 cells). (**g**) *In situ* hybridization in the VTA for *Th* and *Ntsr1*. Scale bars = 150 µm (greyscale images) and 25 µm (zoom). (**h**) Quantification of *Th* and *Ntsr1* expressing neurons of the VTA. (**i**) Example traces and quantification of currents evoked by bath NTS in VTA-DA (black) and VTA-GABA neurons (green); (scale bar=50 pA, 20 s, n=4-10 cells/data point from N=2-3 mice). Data are presented as mean ± standard error of the mean (SEM). (**b**,**c and h**, see Extended Data Table 1 for cell counts).

Within the VTA, *Ntsr1* is reportedly expressed almost exclusively in DA neurons^17-19^; we confirmed this with *in situ* hybridization for *Ntsr1* and *Th* (Fig. 1g,h). Bath application of Nts evokes a depolarizing current in VTA neurons^20^, and we found that this effect is selective for VTA-DA, but not VTA-GABA neurons (Fig. 1i). Consistent with the specificity of Ntsr1 in mediating the observed effects, we found that the expression of the alternate Nts receptor encoding gene, *Ntsr2*, is almost exclusively in glial cells and not neurons (Extended Data Fig. 1c-d).

We next devised a strategy that allows us to inactivate GABA release from LH-Nts neurons through Cre-dependent CRISPR/SaCas9 mutagenesis of *Vgat* and simultaneously inactivate Nts signaling in VTA-DA neurons through Flp-dependent mutagenesis of *Ntsr1*. We previously validated mutagenesis at the Vgat locus using AAV1-FLEX-SaCas9-U6-sgVgat^2^; we confirmed that this virus effectively reduces GABAergic transmission from LH-Nts to VTA-GABA neurons compared to control (AAV1-FLEX-SaCas9-U6-sgRosa26^2^; Fig. 2a). We next generated a Flp-dependent CRISPR virus containing a single guide RNA targeting *Ntsr1* (AAV-FLEXfrt-SaCas9-U6-sgNtsr1); we confirmed effective mutagenesis in *Th*-Flp neurons through DNA sequencing (Extended Data Fig. 2a-d). We also found that Nts-induced depolarization of VTA-DA neurons in slice is blocked by CRISPR/SaCas9 mutagenesis of *Ntsr1* (Fig. 2b), without affecting glutamatergic or GABAergic synaptic inputs (Extended Data Fig 2e-j).

**Figure 2.**
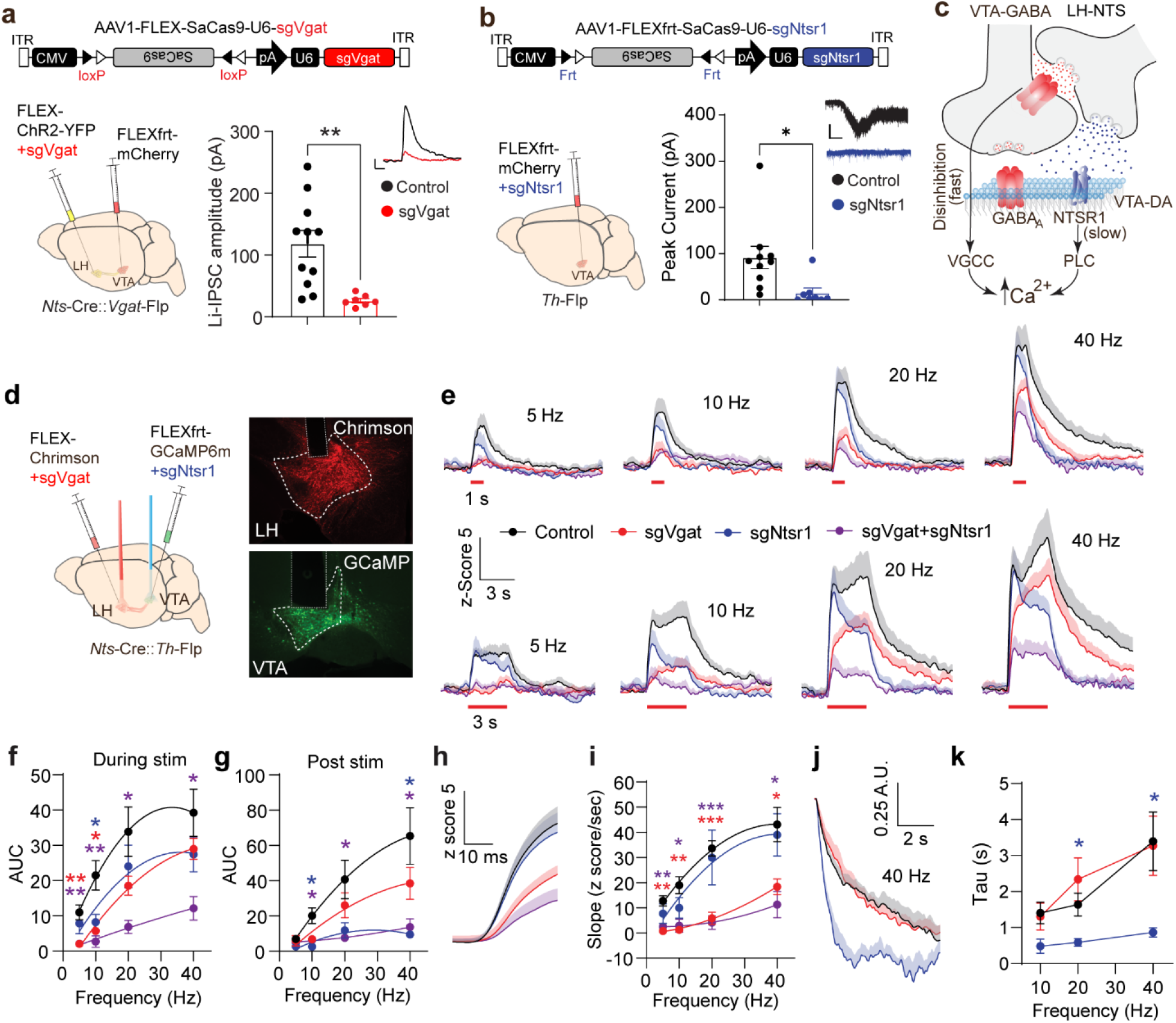
LH-Nts evoked calcium signals in the VTA are regulated by GABA release and Ntsr1 signaling. (**a**) Schematic of Cre-dependent CRISPR virus targeting Vgat (top); schematic of viral injections (bottom left); and quantification and example traces of Li-IPSCs recorded in VTA-GABA neurons with control CRISPR or sgVgat expression in LH-NTS neurons (bottom right; control, n=11 cells from N=3 mice; sgVgat n=7 cells from N=3 mice; **P<0.01, scale bar: 25 pA, 10 ms). **(b)** Schematic of Flp-dependent CRISPR virus targeting *Ntsr1* (top); schematic of viral injections (bottom left); and quantification and example traces of currents evoked by bath NTS (bottom right; control, n=10 cells from N=3 mice; sgNtsr1 n=8 cells from N=2 mice; *P<0.05, scale bar: 20 s, 50 pA). (**c**) Proposed model of calcium signals in VTA-DA neurons generated by GABA disinhibition and Ntsr1 signaling. (Note: the axo-axonic connection is presented for simplicity and is not meant to reflect the only type of synapse involved in co-release). (**d**) Schematic of viral injection and fiber implantation (left) and expression of Chrimson-tdTomato and GCaMP6m in the LH and VTA, respectively, with representative optical fiber placements for stimulation and imaging (right). (**e**) Average GCaMP6m fluorescence (Z-score) following 1 s (top) or 3 s (bottom) red light stimulation at different frequencies. (**f-g**) Average area under the curve (AUC) of the Z-scored GCaMP6m fluorescence during (**f**) and for 10 s following (**g**) red light stimulation at different frequencies (3 s stim). (**h)** Average GCaMP6m fluorescence (Z-score) at the onset of 3 s, 40 Hz optical stimulation. **(i)** Average slope of the initial rise in GCaMP6m fluorescence at different stimulus frequencies (3 s stim). **(j)** Average GCaMP6m fluorescence decay following termination of 3 s stimulation, normalized to peak. (**k**) Average decay time constant (tau) of GCaMP6m fluorescence at different stimulus frequencies for 3 s stimulation. Note: Analysis of 5 Hz was omitted due to lack of signal in sgVgat mice, and sgVgat+sgNtsr1 mice were excluded from analysis due to the low amplitude of the evoked fluorescence. (**e-k**, control N=8, sgVgat N=7, sgNtsr1 N=6, sgVgat+sgNtsr1 N=5; *P<0.05, **P<0.01, ***P<0.001, red asterisk sgVgat vs. control; blue asterisk sgNtsr1 vs. control; purple asterisk sgVgat+sgNtsr1 vs. control). Data are presented as mean ± SEM.

In our proposed model (Fig. 2c and Extended Data Fig. 3a), GABA release from LH-Nts neurons onto VTA-GABA neurons disinhibits VTA-DA neurons, depolarizing these cells and activating voltage-gated calcium channels (VGCC)^21^. Activation of the Gq-coupled Ntsr1 on these cells by Nts release from LH-Nts neurons will also increase intracellular calcium on a slower timescale through phospholipase C (PLC) signaling^22^. These changes in intracellular calcium can be monitored by expression of the genetically encoded calcium sensor GCaMP6m^23^. To test whether activation of LH-Nts inputs to the VTA increases intracellular calcium in dopamine neurons in a GABA- and Ntsr1-dependent manner, we expressed the red light-activated opsin Chrimson^24^ in LH-Nts neurons (AAV1-FLEX-Chrimson-Tdtomato), mutated the *Vgat* locus in LH-Nts neurons alone, mutated *Ntsr1* in VTA-DA neurons alone, or mutated *Vgat* in LH-Nts neurons and *Ntsr1* in VTA-DA neurons together, while expressing GCaMP6m in VTA-DA neurons in all conditions (AAV1-FLEXfrt-GCaMP6m) (Fig. 2d; Extended Data Fig. 3b-e and Extended Data Fig. 4).

Optical stimulation of LH-Nts neurons for 1 or 3 s reliably evoked calcium signals in VTA-DA neurons during the stimulation period (Fig. 2e-f; Extended Data Fig. 5a-c). We also observed a prolonged calcium signal following stimulus offset that increased with stimulus frequency and duration (Fig. 2g; Extended Data Fig. 5d). Inactivation of *Ntsr1* in VTA-DA neurons or *Vgat* in LH-Nts neurons each reduced evoked calcium signals, but with different kinetics and frequency dependence. Targeting *Vgat* had the most pronounced effect during the stimulation period at low frequency and duration, while targeting *Ntsr1* strongly reduced the post-stimulation calcium signal, particularly at high frequencies (Fig. 2e-g; Extended Data Fig. 5a-f). Simultaneous mutagenesis of both genes in their respective cell types further reduced evoked calcium signals (Fig. 2e-g; Extended Data Fig. 5a-f).

During LH-Nts neuron stimulation at 10 Hz or higher we detected two distinct peaks in the calcium signal in VTA-DA neurons that were most prominent with the 3 s stimulation (Fig. 2e, Extended Data Fig. 5a-b). Further analysis revealed that the initial peak was reduced by inactivation of *Vgat* in LH-Nts neurons and the second peak was reduced relative to the first peak by inactivation of *Ntsr1* in VTA-DA neurons (Extended Data Fig. 5g-l). To further resolve the fast kinetics of the calcium signal, we analyzed the initial rise time during the 3 s stimulation. Mutagenesis of *Vgat* in LH-Nts neurons significantly attenuated this initial rise but *Ntsr1* mutagenesis did not affect these kinetics (Fig. 2h-i). We observed identical results when we analyzed the initial rise of the 1 s stimulation (Extended Data Fig. 6a-b). Next, we assessed the decay of the calcium signal following termination of the LH-Nts neuron stimulation. Inactivation of *Ntsr1* in VTA-DA neurons significantly increased the rate of decay (decreased *tau*) following both 1 and 3s stimulation, while inactivation of *Vgat* in LH-Nts neurons did not significantly affect decay (Fig. 2j-k and Extended Data Fig. 6c-d). These results demonstrate that GABA release from LH-Nts neurons and Ntsr1 activation regulate calcium signals in VTA-DA neurons on different time scales in a frequency-dependent manner. GABA is the primary regulator of the calcium signal at low frequencies and at the onset of high frequency stimulation, while Ntsr1 mediates a slow onset, long lasting calcium signal detectable at frequencies of 10 Hz and above, and has a larger impact than GABA on the total calcium signal at high frequencies and long durations.

We next asked how the fast transmitter and peptide components of this circuit might contribute to dopamine neuron activation during reinforcement learning. We first used photometry to record calcium signals in LH-Nts neurons while mice were trained on an operant paradigm in which one press on an active lever led to a delayed compound conditioned stimulus (CS, tone+light) followed by a food reward (Fig. 3a-b and Extended Data Fig. 7a). We observed a small decrease in fluorescence during the CS presentation on day 1, and a large increase in fluorescence upon head entry into the food hopper to retrieve the reward on both day 1 and day 5 of training (Fig. 3d), consistent with a proposed role for LH-Nts neurons in feeding behavior^25^.

**Figure 3.**
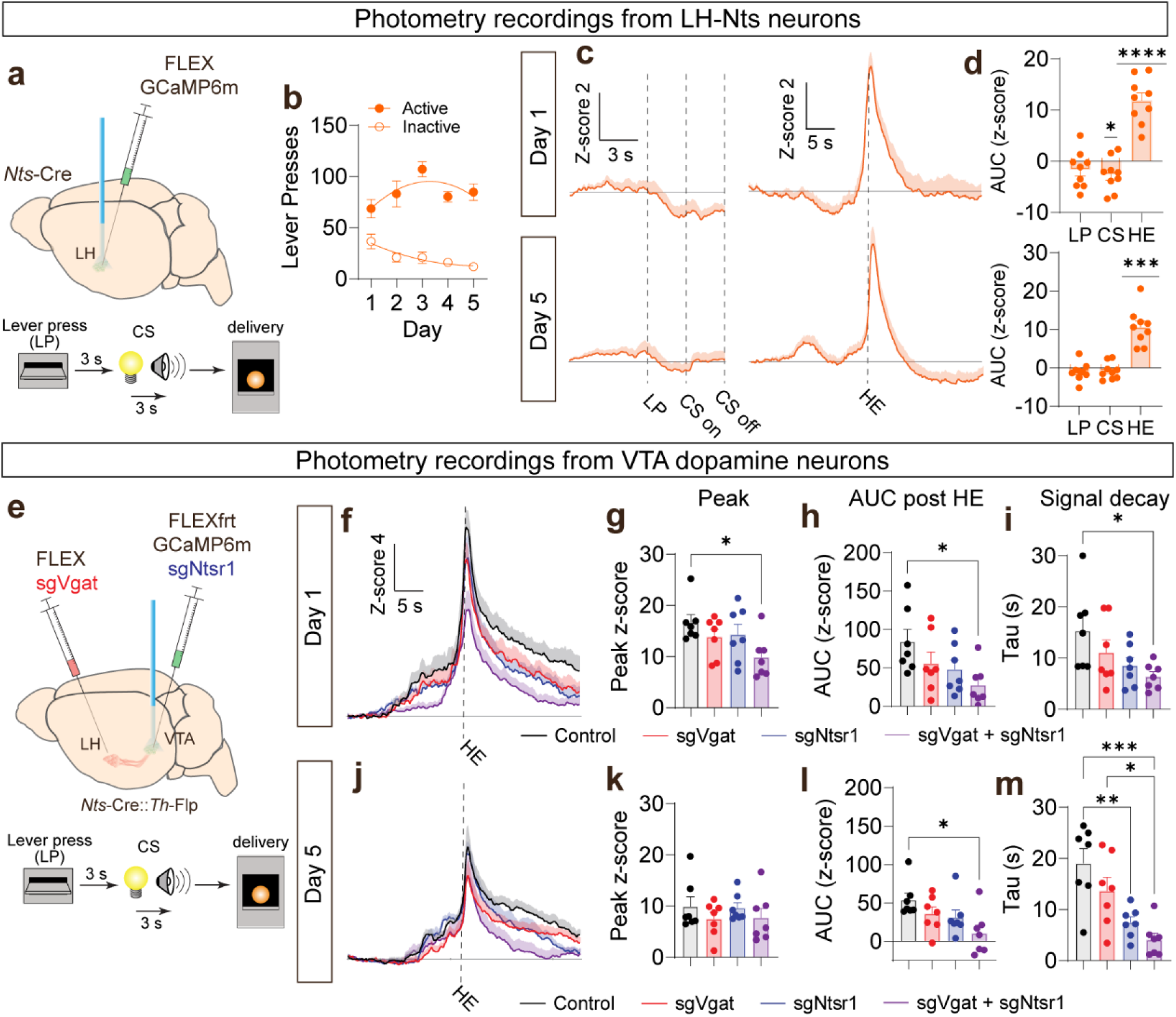
LH-Nts neurons contribute to activation of VTA-DA neurons during reward retrieval. **(a)** Schematic of viral injection and fiber implantation to record from LH-Nts neurons (top), schematic of operant paradigm (bottom). **(b)** Active and inactive lever presses across 5 days of training. **(c)** Average GCaMP6m fluorescence (Z-score) in LH-Nts neurons on day 1 (top) or day 5 (bottom) of operant conditioning. Left: response to lever press (LP) and cue (CS) presentation. Right: response to first head entry (HE) into the food hopper following reward delivery. **(d)** AUC for the 3 s following the LP, the 3 s during the cue delivery, or the 3 s following HE (N=9 mice, 1-sample t-test to determine significant difference from zero: *P<0.05, ***P<0.001, ****P<0.0001). **(e)** Schematic of viral injection and fiber implantation to record from VTA-DA neurons (top), schematic of operant paradigm (bottom). **(f)** Average GCaMP6m fluorescence (Z-score) on day 1 of conditioning, aligned to first HE following reward delivery. **(g)** Peak Z-score, **(h)** AUC from 5 to 20 s following HE, and **(i)** Decay time constant (Tau) following HE on day 1 of conditioning. **(j)** Average GCaMP6m fluorescence (Z-score) on day 5 of conditioning, aligned to first HE. **(k)** Peak Z-score, **(l)** AUC from 5 to 20 s following HE, and **(m)** Decay time constant (Tau) following HE, on day 5 of conditioning. (N=7 mice per group. *P<0.05, **P<0.01, ***P<0.001 for indicated comparisons.) Data are presented as mean ± SEM.

Next we recorded calcium signals in dopamine neurons following CRISPR targeting of *Vgat* in LH-Nts neurons, *Ntsr1* in VTA-DA neurons, or both, while mice were trained on the same operant paradigm (Fig. 3e and Extended Data Fig. 7b). We observed large calcium signals time-locked to reward retrieval head entries in VTA-DA neurons on day 1 of training (Figure 3f). Combined CRISPR targeting of *Vgat* and *Ntsr1* reduced the peak of this calcium signal, reduced the AUC during the prolonged decay of the signal following the head entry, and increased the rate of decay compared to control (Fig. 3g-i). Control and sgVgat animals showed a smaller head entry response on day 5 of training compared to day 1 (Fig. 3j-k and Extended Data Fig.8a-d), consistent with models of reward prediction error^26^. The AUC during the post-head entry period remained reduced in sgVgat+sgNtsr1 animals on day 5 (Fig. 3l), and the rate of decay was significantly faster in sgNtsr1 alone and sgVgat+sgNtsr1 animals compared to control (Fig. 3m), consistent with Ntsr1 mediating the slow calcium signal.

Calcium signals in dopamine neurons in response to the lever press and cue were not different between groups (Extended data Fig. 8e-l), and we did not observe a significant difference in the number of earned rewards (Extended data Fig. 8m-q). However, across all trials the mean latency to make a head entry into the food hopper following reward delivery was significantly slower in the final quarter of the session on day 5 in double mutant mice compared to controls (Extended data Fig. 8r-s), potentially indicating increased satiety or reduced motivation to retrieve the pellet. Intriguingly, in a separate cohort of mice we observed that sgVgat+sgNtsr1 mice exhibit a significant loss of body weight compared to controls 5 weeks post-viral injection (Extended data Fig. 8t). Similar results were found after silencing LH-Nts neurons with Cre-dependent expression of the tetanus toxin light chain (AAV1-FLEX-GFP-TeTox; Extended data Fig. 8t).

Activation of LH to VTA GABA projections is strongly reinforcing^8,14^. To determine whether activation of LH-Nts inputs to the VTA is sufficient to drive reinforcement and if either Nts signaling in the VTA or GABA release from LH neurons is required for these reinforcing effects, we inactivated *Ntsr1* in VTA-DA neurons or *Vgat* in LH-Nts neurons, as above, expressed Chrimson-tdTomato in LH-Nts neurons and stimulated the terminals of these cells in the VTA during a real-time place preference assay (Fig. 4a). Stimulation of LH-Nts inputs (20 Hz) promoted a preference for the light-paired chamber and neither inactivation of *Ntsr1* nor *Vgat* affected this preference (Fig. 4b; Extended Data Fig. 9a-b); however, inactivation of both genes simultaneously blocked this behavioral response (Fig. 4a; Extended Data Fig. 9a-b).

**Figure 4.**
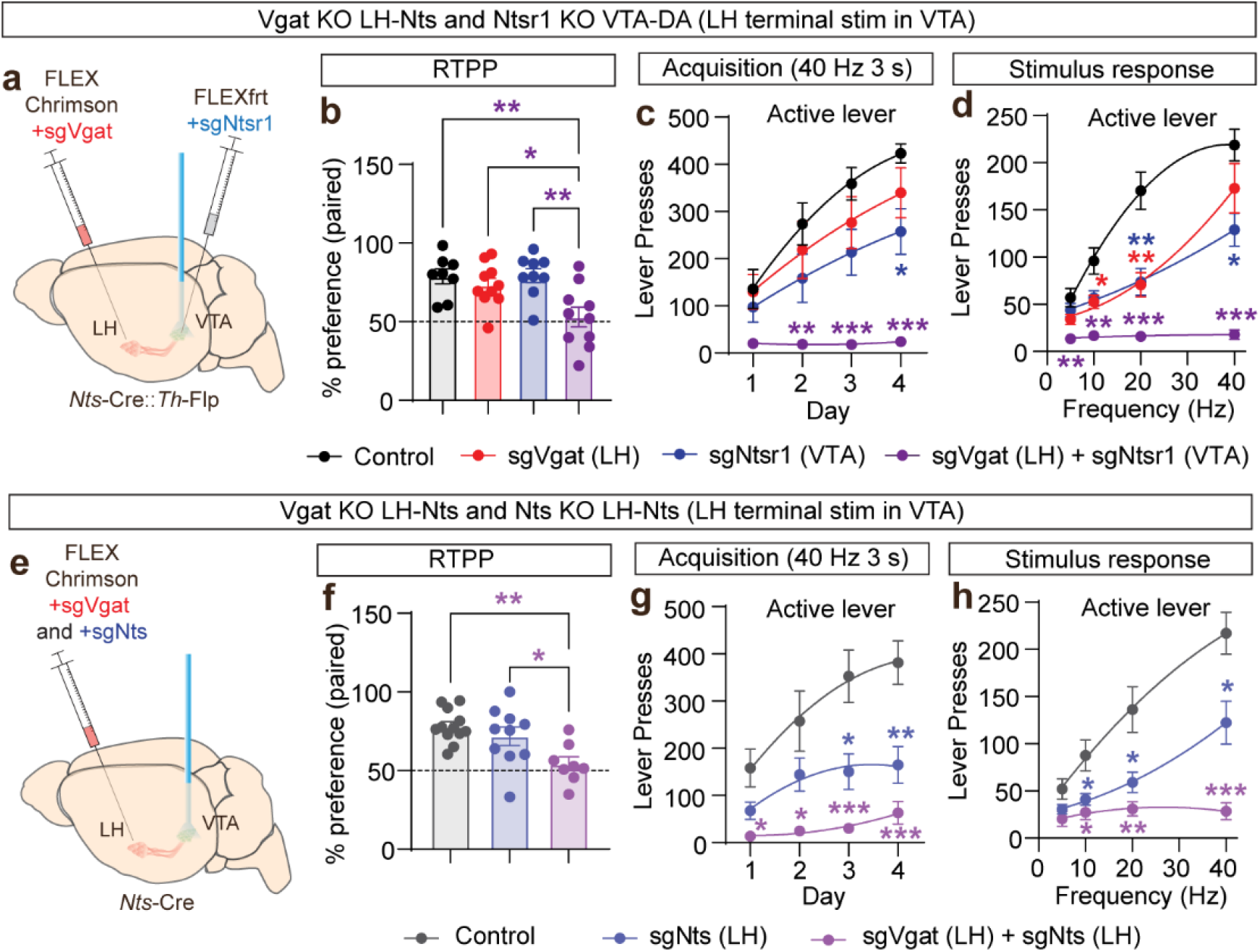
Coordinated actions of LH-Nts and GABA co-release in behavioral reinforcement. (**a**) Schematic of viral injections and stimulating fiber placement. **(b)** Average percent time in light-paired (20 Hz) chamber during final 10 min of RTPP (**P<0.01; *P<0.05; control N=8, sgVgat N=,10 sgNtsr1 N=9, sgVgat+sgNtsr1 N=10). (**c**) Average active lever press responses (60 min session) for 40 Hz, 3 s optical stimulation of LH-Nts inputs to the VTA. (**d**) Average active lever presses (30 min session) for optical stimulation of LH-Nts inputs to the VTA at different frequencies (**c**,**d** ***P<0.001; **P<0.01; *P<0.05; red asterisk sgVgat vs. control; blue asterisk sgNtsr1 vs. control; purple asterisk sgVgat+sgNtsr1 vs. control; control N=9, sgVgat N=,10 sgNtsr1 N=8, sgVgat+sgNtsr1 N=10). (**e**) Schematic of viral injections and stimulating fiber placement. (**f**) Average percent time in light-paired (20 Hz) chamber during final 10 min of RTPP (**P<0.01; *P<0.05; control N=12; sgNts=10; sgNts+sgVgat N=8). (**g**) Average active lever press responses (60 min session) for 40 Hz, 3 s optical stimulation of LH-Nts inputs to the VTA. (**h**) Average active lever presses (30 min session) for optical stimulation of LH-Nts inputs to the VTA at different frequencies (**g**,**h** ***P<0.001; **P<0.01; *P<0.05; blue asterisk sgNtsr1 vs. control; purple asterisk sgVgat+sgNtsr1 vs. control; control N=9; sgNts N=11; sgVgat+sgNts N=7). Data are presented as mean ± SEM.

To better resolve the interaction between Ntsr1 signaling in VTA-DA neurons and GABA release from LH-Nts neurons, as well as the frequency dependence of this interaction, we trained mice on a fixed-ratio schedule 1 (FR1) operant (lever press) reinforcement task for optical activation of LH-Nts terminals in the VTA. Activation of LH-Nts inputs to the VTA (40 Hz, 3 s) resulted in robust lever pressing in control mice (Fig. 4c; Extended data Fig. 9c). Mice with inactivation of *Vgat* in LH-Nts neurons or *Ntsr1* in VTA-DA neurons both learned to lever press at this stimulation frequency (Fig. 4c; Extended Data Fig. 9d-e); however, inactivation of *Ntsr1* significantly reduced lever pressing relative to controls (Fig. 4c). Mutagenesis of both *Ntsr1* in VTA-DA neurons and *Vgat* in LH-Nts neurons prevented the acquisition of this response (Fig. 4c; Extended Data Fig. 9f). Following FR1 training for 40 Hz stimulation, we assayed mice in a frequency-response assay. Mice were allowed to lever press for optical activation of LH-Nts inputs at 5, 10, 20, or 40 Hz. Control mice displayed a robust frequency-response curve for activation of LH-Nts inputs to the VTA (Fig. 4d; Extended Data Fig. 9g) that was significantly attenuated following inactivation of either *Vgat* or *Ntsr1* (Fig. 4d; Extended Data Fig. 9h-i) and was completely blocked by mutagenesis of both genes in their respective cell types (Fig. 4d; Extended Data Fig. 9j).

Our results demonstrate that GABA release from LH-Nts neurons and Ntsr1 activation in VTA-DA neurons are required for optimal behavioral reinforcement following activation of this circuit connection but do not directly demonstrate that Nts and GABA co-release from LH-Nts neurons mediates this effect. To address this, we inactivated the *Nts* gene (AAV1-FLEX-SaCas9-U6-sgNts; Extended Data Fig. 10a-d) or both *Nts* and *Vgat* in LH-Nts neurons and repeated the experiments described above. Like mutagenesis of *Ntsr1* in VTA-DA neurons, inactivation of *Nts* did not affect RTPP, but inactivation of both *Nts* and *Vgat* did prevent this preference (Fig. 4e-f; Extended Data Fig. 10e-f). Also similar to inactivation of *Ntsr1*, mutagenesis of *Nts* significantly attenuated the acquisition of operant stimulation of LH-Nts terminals in the VTA and the frequency-response profile, which was further reduced by inactivation of both *Vgat* and *Nts* (Fig. 4g-h; Extended Data Fig. 10g-l).

These results resolve the contribution of Nts and GABA co-release from LH-Nts neurons to behavioral reinforcement through projections to the VTA. We describe a circuit mechanism whereby the co-release of a neurotransmitter and neuropeptide with ostensibly opposing effects on cell physiology (depolarization versus hyperpolarization) can work coordinately to promote output from a circuit node by acting at different time scales on distinct cell types. The activation of LH-Nts neurons to food reward, combined with the reduction of calcium signals in dopamine neurons in response to the same reward following inactivation of *Vgat* and *Ntsr1*, provides evidence that both LH-GABA and Nts signaling in dopamine neurons contribute to the reinforcing effects of food. This is consistent with our optogenetics experiments demonstrating that both signals coordinately contribute to reinforcement.

These findings provide new insights into the circuit organization of a critical input to the dopamine system. Given the large number of GABAergic inputs to the VTA that disinhibit dopamine neurons and are likely to co-release excitatory peptides^8,27^, as well as the enrichment of many peptide receptors on dopamine neurons^28^, this may prove to be a common mechanism for regulation of the dopamine system. The experimental approach described here for *in vivo* isolation of circuit components can be easily adapted going forward to investigate other peptidergic inputs to the VTA, as well as peptidergic circuits throughout the brain to address these fundamental questions (also see Supplementary Discussion).

## Acknowledgments

Supported by National Institutes of Health grants R21 MH121774 (MES), R01 DA054924 (MES), and R01 DA044315 (LSZ). This work was also supported by the University of Washington Center of Excellence in Opioid Addiction Research (P30 DA048736). We thank Dr. James Allen, Dr. Avery Hunker, and Dasha Krayushkina for assistance with viral production, and Dr. Scott Ng-Evans for assistance with photometry analysis. We also thank members of the Zweifel lab for their thoughtful discussion.

## Author contributions

M.E.S. and L.S.Z conceptualized the study and designed experiments. M.E.S. and J.X.Y. performed all experiments and collected data. M.E.S., J.X.Y., and L.S.Z. analyzed data and wrote the paper.

## Data availability

All data associated with this study will be made available by the corresponding author upon reasonable request.

## References

1 Bartfai, T., Iverfeldt, K., Fisone, G. & Serfozo, P. Regulation of the release of coexisting neurotransmitters. Annu Rev Pharmacol Toxicol 28, 285–310, doi:10.1146/annurev.pa.28.040188.001441 (1988).

2 Hunker, A. C. et al. Conditional Single Vector CRISPR/SaCas9 Viruses for Efficient Mutagenesis in the Adult Mouse Nervous System. Cell Rep 30, 4303–4316 e4306, doi:10.1016/j.celrep.2020.02.092 (2020).

3 Salio, C., Lossi, L., Ferrini, F. & Merighi, A. Neuropeptides as synaptic transmitters. Cell Tissue Res 326, 583–598, doi:10.1007/s00441-006-0268-3 (2006).

4 van den Pol, A. N. Neuropeptide transmission in brain circuits. Neuron 76, 98–115, doi:10.1016/j.neuron.2012.09.014 (2012).

5 Tritsch, N. X., Granger, A. J. & Sabatini, B. L. Mechanisms and functions of GABA co-release. Nat Rev Neurosci 17, 139–145, doi:10.1038/nrn.2015.21 (2016).

6 Brown, J. A. et al. Distinct Subsets of Lateral Hypothalamic Neurotensin Neurons are Activated by Leptin or Dehydration. Sci Rep 9, 1873, doi:10.1038/s41598-018-38143-9 (2019).

7 Mickelsen, L. E. et al. Single-cell transcriptomic analysis of the lateral hypothalamic area reveals molecularly distinct populations of inhibitory and excitatory neurons. Nat Neurosci 22, 642–656, doi:10.1038/s41593-019-0349-8 (2019).

8 Soden, M. E. et al. Anatomic resolution of neurotransmitter-specific projections to the VTA reveals diversity of GABAergic inputs. Nat Neurosci 23, 968–980, doi:10.1038/s41593-020-0657-z (2020).

9 Wise, R. A. Dopamine, learning and motivation. Nat Rev Neurosci 5, 483–494, doi:10.1038/nrn1406 (2004).

10 Olds, J. & Milner, P. Positive reinforcement produced by electrical stimulation of septal area and other regions of rat brain. J Comp Physiol Psychol 47, 419–427, doi:10.1037/h0058775 (1954).

11 Olds, J., Travis, R. P. & Schwing, R. C. Topographic organization of hypothalamic self-stimulation functions. J Comp Physiol Psychol 53, 23–32, doi:10.1037/h0039776 (1960).

12 Jennings, J. H. et al. Visualizing hypothalamic network dynamics for appetitive and consummatory behaviors. Cell 160, 516–527, doi:10.1016/j.cell.2014.12.026 (2015).

13 Kempadoo, K. A. et al. Hypothalamic neurotensin projections promote reward by enhancing glutamate transmission in the VTA. J Neurosci 33, 7618–7626, doi:10.1523/JNEUROSCI.2588-12.2013 (2013).

14 Nieh, E. H. et al. Inhibitory Input from the Lateral Hypothalamus to the Ventral Tegmental Area Disinhibits Dopamine Neurons and Promotes Behavioral Activation. Neuron 90, 1286–1298, doi:10.1016/j.neuron.2016.04.035 (2016).

15 Patterson, C. M. et al. Ventral tegmental area neurotensin signaling links the lateral hypothalamus to locomotor activity and striatal dopamine efflux in male mice. Endocrinology 156, 1692–1700, doi:10.1210/en.2014-1986 (2015).

16 Boyden, E. S., Zhang, F., Bamberg, E., Nagel, G. & Deisseroth, K. Millisecond-timescale, genetically targeted optical control of neural activity. Nat Neurosci 8, 1263–1268, doi:10.1038/nn1525 (2005).

17 Szigethy, E. & Beaudet, A. Correspondence between high affinity 125I-neurotensin binding sites and dopaminergic neurons in the rat substantia nigra and ventral tegmental area: a combined radioautographic and immunohistochemical light microscopic study. J Comp Neurol 279, 128–137, doi:10.1002/cne.902790111 (1989).

18 Opland, D. et al. Loss of neurotensin receptor-1 disrupts the control of the mesolimbic dopamine system by leptin and promotes hedonic feeding and obesity. Mol Metab 2, 423–434, doi:10.1016/j.molmet.2013.07.008 (2013).

19 Woodworth, H. L., Perez-Bonilla, P. A., Beekly, B. G., Lewis, T. J. & Leinninger, G. M. Identification of Neurotensin Receptor Expressing Cells in the Ventral Tegmental Area across the Lifespan. eNeuro 5, doi:10.1523/ENEURO.0191-17.2018 (2018).

20 Jiang, Z. G., Pessia, M. & North, R. A. Neurotensin excitation of rat ventral tegmental neurones. J Physiol 474, 119–129, doi:10.1113/jphysiol.1994.sp020007 (1994).

21 Catterall, W. A. Voltage-gated calcium channels. Cold Spring Harb Perspect Biol 3, a003947, doi:10.1101/cshperspect.a003947 (2011).

22 Snider, R. M., Forray, C., Pfenning, M. & Richelson, E. Neurotensin stimulates inositol phospholipid metabolism and calcium mobilization in murine neuroblastoma clone N1E-115. J Neurochem 47, 1214–1218, doi:10.1111/j.1471-4159.1986.tb00742.x (1986).

23 Chen, T. W. et al. Ultrasensitive fluorescent proteins for imaging neuronal activity. Nature 499, 295–300, doi:10.1038/nature12354 (2013).

24 Klapoetke, N. C. et al. Independent optical excitation of distinct neural populations. Nat Methods 11, 338–346, doi:10.1038/nmeth.2836 (2014).

25 Schroeder, L. E. & Leinninger, G. M. Role of central neurotensin in regulating feeding: Implications for the development and treatment of body weight disorders. Biochim Biophys Acta Mol Basis Dis 1864, 900–916, doi:10.1016/j.bbadis.2017.12.036 (2018).

26 Bromberg-Martin, E. S., Matsumoto, M. & Hikosaka, O. Dopamine in motivational control: rewarding, aversive, and alerting. Neuron 68, 815–834, doi:10.1016/j.neuron.2010.11.022 (2010).

27 Soden, M. E. et al. Distinct Encoding of Reward and Aversion by Peptidergic BNST Inputs to the VTA. Front Neural Circuits 16, 918839, doi:10.3389/fncir.2022.918839 (2022).

28 Chung, A. S., Miller, S. M., Sun, Y., Xu, X. & Zweifel, L. S. Sexual congruency in the connectome and translatome of VTA dopamine neurons. Sci Rep 7, 11120, doi:10.1038/s41598-017-11478-5 (2017).

29 Nusbaum, M. P., Blitz, D. M. & Marder, E. Functional consequences of neuropeptide and small-molecule co-transmission. Nat Rev Neurosci 18, 389–403, doi:10.1038/nrn.2017.56 (2017).

30 Schone, C. & Burdakov, D. Glutamate and GABA as rapid effectors of hypothalamic “peptidergic” neurons. Front Behav Neurosci 6, 81, doi:10.3389/fnbeh.2012.00081 (2012).

31 Schone, C., Apergis-Schoute, J., Sakurai, T., Adamantidis, A. & Burdakov, D. Coreleased orexin and glutamate evoke nonredundant spike outputs and computations in histamine neurons. Cell Rep 7, 697–704, doi:10.1016/j.celrep.2014.03.055 (2014).

32 Li, Y. & van den Pol, A. N. Differential target-dependent actions of coexpressed inhibitory dynorphin and excitatory hypocretin/orexin neuropeptides. J Neurosci 26, 13037–13047, doi:10.1523/JNEUROSCI.3380-06.2006 (2006).

33 Apergis-Schoute, J. et al. Optogenetic evidence for inhibitory signaling from orexin to MCH neurons via local microcircuits. J Neurosci 35, 5435–5441, doi:10.1523/JNEUROSCI.5269-14.2015 (2015).

34 Muschamp, J. W. et al. Hypocretin (orexin) facilitates reward by attenuating the antireward effects of its cotransmitter dynorphin in ventral tegmental area. Proc Natl Acad Sci U S A 111, E1648–1655, doi:10.1073/pnas.1315542111 (2014).

35 McHenry, J. A. et al. Hormonal gain control of a medial preoptic area social reward circuit. Nat Neurosci 20, 449–458, doi:10.1038/nn.4487 (2017).

36 Dedic, N. et al. Chronic CRH depletion from GABAergic, long-range projection neurons in the extended amygdala reduces dopamine release and increases anxiety. Nat Neurosci 21, 803–807, doi:10.1038/s41593-018-0151-z (2018).

37 Perez-Bonilla, P., Santiago-Colon, K. & Leinninger, G. M. Lateral hypothalamic area neuropeptides modulate ventral tegmental area dopamine neurons and feeding. Physiol Behav 223, 112986, doi:10.1016/j.physbeh.2020.112986 (2020).

38 Heymann, G. et al. Synergy of Distinct Dopamine Projection Populations in Behavioral Reinforcement. Neuron 105, 909–920 e905, doi:10.1016/j.neuron.2019.11.024 (2020).

39 Paladini, C. A. & Tepper, J. M. GABA(A) and GABA(B) antagonists differentially affect the firing pattern of substantia nigra dopaminergic neurons in vivo. Synapse 32, 165–176, doi:10.1002/(SICI)1098-2396(19990601)32:3<165::AID-SYN3>3.0.CO;2-N (1999).

40 Sotty, F. et al. Differential effects of neurotensin on dopamine release in the caudal and rostral nucleus accumbens: a combined in vivo electrochemical and electrophysiological study. Neuroscience 85, 1173–1182, doi:10.1016/s0306-4522(97)00691-x (1998).

41 Destexhe, A. & Marder, E. Plasticity in single neuron and circuit computations. Nature 431, 789–795, doi:10.1038/nature03011 (2004).

42 Hawkins, M. F. Aphagia in the rat following microinjection of neurotensin into the ventral tegmental area. Life Sci 38, 2383–2388, doi:10.1016/0024-3205(86)90606-5 (1986).

43 Woodworth, H. L. et al. Lateral Hypothalamic Neurotensin Neurons Orchestrate Dual Weight Loss Behaviors via Distinct Mechanisms. Cell Rep 21, 3116–3128, doi:10.1016/j.celrep.2017.11.068 (2017).

44 Remaury, A. et al. Targeted inactivation of the neurotensin type 1 receptor reveals its role in body temperature control and feeding behavior but not in analgesia. Brain Res 953, 63–72, doi:10.1016/s0006-8993(02)03271-7 (2002).

45 de Jong, J. W. et al. A Neural Circuit Mechanism for Encoding Aversive Stimuli in the Mesolimbic Dopamine System. Neuron 101, 133–151 e137, doi:10.1016/j.neuron.2018.11.005 (2019).

46 Paxinos, G. & Franklin, K. B. J. The mouse brain in stereotaxic coordinates. Compact 2nd edn, (Elsevier Academic Press, 2004).

47 Poulin, J. F. et al. Mapping projections of molecularly defined dopamine neuron subtypes using intersectional genetic approaches. Nat Neurosci 21, 1260–1271, doi:10.1038/s41593-018-0203-4 (2018).

48 Gore, B. B., Soden, M. E. & Zweifel, L. S. Manipulating gene expression in projection-specific neuronal populations using combinatorial viral approaches. Curr Protoc Neurosci 65, 4 35 31–20, doi:10.1002/0471142301.ns0435s65 (2013).

49 Ting, J. T., Daigle, T. L., Chen, Q. & Feng, G. Acute brain slice methods for adult and aging animals: application of targeted patch clamp analysis and optogenetics. Methods Mol Biol 1183, 221–242, doi:10.1007/978-1-4939-1096-0_14 (2014).

50 Hunker, A. C. & Zweifel, L. S. Protocol to Design, Clone, and Validate sgRNAs for In Vivo Reverse Genetic Studies. STAR Protoc 1, doi:10.1016/j.xpro.2020.100070 (2020).

51 Brinkman, E. K., Chen, T., Amendola, M. & van Steensel, B. Easy quantitative assessment of genome editing by sequence trace decomposition. Nucleic Acids Res 42, e168, doi:10.1093/nar/gku936 (2014).

