## Supplementary material for "A circuit mechanism for the coordinated actions of opposing neuropeptide and neurotransmitter signals"

### Methods

**Mice:** All procedures were approved and conducted in accordance with the guidelines of the Institutional Animal Care and Use Committee of the University of Washington. Mice were group-housed on a 12-hour light/dark cycle with ad libitum food and water. Approximately equal numbers of male and female mice were used for all experiments. Nts-Cre mice (Jax Stock Number 017525) and Vgat-Flp mice (Jax Stock Number 029591) are available from Jackson Labs. Th-Flp mice[47] were a generous gift from Dr. Rajeshwar Awatramani.

**Viruses:** All AAV1 viruses were produced in-house with titers of 1-3 $x10^{12}$ particles per mL as described[48].

**Surgery:** Mice were anesthetized with isoflurane before and during viral injection and fiber implantation. Mice recovered for at least 4 weeks prior to experimentation. For slice electrophysiology, mice were injected at approximately 5-6 weeks of age. For all other experiments, mice were injected at 8-12 weeks of age. LH coordinates were M-L: 1.0, A-P: -1.25, D-V: -5.0. VTA coordinates were M-L: 0.5, A-P: -3.25, D-V:-4.25. Values are in mm, relative to bregma. A-P values were adjusted for bregma-lambda distance using a correction factor of 4.21 mm. For Z values the syringe was lowered 0.5 mm past the indicated depth and raised up at the start of the injection. Injection volume was 500 nl. Injections for slice electrophysiology and behavioral photometry were bilateral; injections for stimulated photometry and optogenetic behavior were unilateral.

Fiber optic cannulas for *in vivo* Chrimson stimulation were manufactured in-house using 1.25 mm ceramic ferrules (Kientec) and 200 µm 0.22 NA fiber (ThorLabs). Fiber optic cannulas for photometry (400 µm fiber, 0.66 NA, 1.25 mm ferrule) were from Doric. For photometry, the imaging cannula was implanted in the VTA at a depth of -4.2 mm from bregma. The stimulating cannula was implanted above the ipsilateral LH at a depth of -4.8 mm and an angle of 5°. For optogenetic behavior experiments, a stimulating cannula was implanted above the VTA at a depth of -4.0 mm from Bregma.

**In Situ Hybridization:** RNAscope (Advanced Cell Diagnostics) *in situ* hybridization was performed according to manufacturer's instructions on 20 μm fresh-frozen coronal brain sections. Version 1 was used to probe for *Nts*, *Cre, Slc32a1*, and *Slc17a6* in the LH, and *Th* and *Flp* in the VTA. Version 2 was used to probe for *Ntsr1*, *Ntsr2*, *Th*, and *Gfap* in the VTA. 20x images were collected using a Leica SP8X confocal microscope and were analyzed using ImageJ software. For quantification of mRNA reduction by sgNts, mice were injected in one hemisphere with sgRosa26 and in the other hemisphere with sgNts. The integrated pixel intensity of all Nts positive neurons in the LH was summed for each hemisection and compared to the opposite hemisection.

**Slice Electrophysiology:** Horizontal brain slices (200 μm) were prepared in an ice slush solution containing (in mM): 92 NMDG, 2.5 KCl, 1.25 $NaH_2PO_4$, 30 $NaHCO_3$, 20 HEPES, 25 glucose, 2 thiourea, 5 Na-ascorbate, 3 Na-pyruvate, 0.5 $CaCl_2$, 10 $MgSO_4$, pH 7.3-7.4[49]. Slices recovered for ≤12 min in the same solution at 32°C and then were transferred to a room temperature solution including (in mM): 92 NaCl, 2.5 KCl, 1.25 $NaH_2PO_4$, 30 $NaHCO_3$, 20 HEPES, 25 glucose, 2 thiourea, 5 Na-ascorbate, 3 Na-pyruvate, 2 $CaCl_2$, 2 $MgSO_4$. Slices recovered for an additional 45 min before recordings were made in ACSF at 32°C continually perfused over slices at a rate of ~2 ml/min and containing (in mM): 126 NaCl, 2.5 KCl, 1.2 NaH2$PO_4$, 1.2 $MgCl_2$ 11 D-glucose, 18 $NaHCO_3$, 2.4 $CaCl_2$. All solutions were continually bubbled with $O_2/CO_2$. Whole-cell recordings were made using an Axopatch 700B amplifier (Molecular Devices) with filtering at 1 kHz using 4-6 MΩ electrodes. For light-evoked PSC and bath Nts recordings electrodes were filled with an internal solution containing (in mM): 130 K-gluconate, 10 HEPES, 5 NaCl, 1 EGTA, 5 Mg-ATP, 0.5 Na-GTP, pH 7.3, 280 mOsm. Light-evoked synaptic transmission was induced with 5 ms light pulses delivered at 0.1 Hz from an optic fiber placed directly in the bath. Light-evoked EPSCs were measured with holding at -70 mV and light-evoked IPSCs were measured with holding at -30 mV. Amplitudes were calculated from an average of at least 10 events. Cells with a detectable event of at least 10 pA were counted as connected. For bath application of NTS, cells were held at -60 mV and NTS (Sigma) was washed on for 2 min. The 1000 nM Th+ group in Fig 1i is the same as the control group in Fig. 2b. Spontaneous EPSCs were recorded using the K-gluconate internal solution described above with holding at -60 mV in the presence of picrotoxin (100 μM). Spontaneous IPSCs were recorded with an internal solution containing (in mM): 135 KCl, 12 NaCl, 0.5 EGTA, 10 HEPES, 2.5 Mg-ATP, 0.25 Na-GTP, pH 7.3, 280 mOsm. sIPSC recordings were made with holding at -60 mV in the presence of kynurenic acid (2 mM). Events were analyzed using Clampfit software (Molecular Devices).

**Design of CRISPR constructs:** Single guide RNAs targeting *Ntsr1* and *Nts* were designed as described[50]. Primers used for cloning sgNtsr1 into AAV1-FLEXfrt-SaCas9-U6 were: forward CACCGCATCGTCTCCAGTCCGAACT and reverse AAACAGTTCGGACTGGAGACGATGC. Primers used for cloning sgNts into AAV1-FLEX-SaCas9-U6 were: forward CACCGACGTTATCAAGGATATCTTC and reverse AAACGAAGATATCCTTGATAACGTC. AAV1-FLEX-SaCas9-U6-sgVgat and AAV1-FLEX-SaCas9-U6-sgRosa26 were previously published[2]. For all CRISPR experiments, control animals were injected with AAV1-FLEX-SaCas9-U6-sgRosa26 (LH) or AAV1-FLEXfrt-SaCas9-U6-sgRosa26 (VTA)[2].

**Validation of mutagenesis:** Validation of mutagenesis was performed using fluorescence activated cell sorting (FACS), whole-genome amplification (WGA), and sequencing as described[2]. Briefly, 3 Th-Flp mice were injected in the VTA with AAV1-FLEXfrt-SaCas9-U6-sgNtsr1 along with AAV-FLEXfrt-EGFP-KASH to label nuclei for sorting. Four weeks following injection tissue punches of the VTA were collected and EGFP-positive nuclei were isolated using FACS. WGA (REPLI-g, Qiagen) was performed according to manufacturer's instructions followed by targeted sequencing of a 200-300 bp region surrounding the intended cut site. Tracking of indels by decomposition (TIDE) analysis[51] was performed to compare sequence chromatograms from EGFP negative and EGFP positive samples to estimate mutation frequency.

**Fiber photometry:** Mice were connected to stimulating and imaging patch cords or the imaging patch cord alone (Doric Lenses) and placed into an operant chamber. The imaging patch cord was photobleached prior to recording. Recordings were made using an RZ5 BioAmp Processor and Synapse software (Tucker Davis Technologies). A 465 nm LED (531-Hz, sinusoidal, Doric Lenses) was used to excite GCaMP6m. LED intensity was measured at the tip of the optic fiber prior to each recording session and set to 30-40 μW. GCaMP6m fluorescence (525 ± 25 nm) was returned through the same patch cord, bandpass filtered, and recorded by the RZ5. A 405 nm LED (211-Hz, sinusoidal) was used to monitor the isosbestic signal. The 531-Hz and 211-Hz signals were extracted by Synapse software at a sampling rate of 1017.25 Hz. For stimulation experiments, red light (640 nm, 5 mW, 5 ms pulses) was delivered through the stimulating patch cord via a laser (LaserGlow) driven by the RZ5. Mice received 5 presentations of each stimulus frequency (5, 10, 20, and 40 Hz) at each duration (1 and 3 s). Stimuli were presented once per minute in a pseudo-random order. Stimulus onsets, lever press, cue, and head entry events were synced to the photometry recording via TTL delivery for offline analysis. A custom Python script was used to extract and analyze the GCaMP signal surrounding each event. A 30 s window was extracted surrounding each red light stimulus and rewarded lever press (10 s prior and 20 s following), and a 40 s window was extracted surrounding the first head entry following reward delivery (20 s prior and 20 s following). The first 4 s of this window was used as a baseline to calculate the Z-score. For stimulation events, Z-scores for the 5 presentations of each stimulus were averaged for each mouse. For behavioral events, Z-scores for the first 20 events in the session were averaged for each mouse.

For stimulation experiments area under the curve during the 1 or 3 s stimulus and 10 s following stimulus termination was calculated relative to the 1 s period prior to stimulation. Peak amplitudes were measured as the maximum value in a 100 ms window surrounding the time of the average initial and final peak measured at 40 Hz in control animals. To quantify the initial rise, a linear regression was performed on the first 20 ms following the stimulus onset. The decay time constant (Tau) was calculated as the time for the signal to decay to 1/e of the final peak amplitude. Maximum values for Tau were capped at 8 s for stimulated photometry and 30 s for behavioral photometry.

**Delayed cue operant task:** Mice were food restricted to 85% of ad libitum body weight. Mice received one 30-minute session of acclimation to the photometry patch cord (Doric Lenses) in the operant chamber (MedAssociates), followed by two pre-training days. On pre-training day 1 mice received 20 non-contingent pellets dispensed with a variable 90 s ITI. On pre-training day 2 the house light was illuminated and both levers were extended. A press on either lever led to extinction of the house light and immediate pellet delivery, followed by a 3 sec ITI. Mice were allowed a maximum of 20 pellets. Next, mice experienced 5 days of delayed cue training, in which both levers were extended but only one lever was active. A press on the active lever led to a 3 s delay, followed by a delivery of a 3 s compound cue (lever light plus 4 KHz tone), followed by pellet delivery. The house light extinguished after each rewarded press and came back on after a 12.5 s ITI to signal the start of a new trial. Training sessions lasted for 1 hour.

**Optogenetics:**

Real-time place preference: On day 1, mice were connected to a patch cable and placed into a two-chambered arena and allowed to explore freely for 10 min. Mice were then assigned a light-paired chamber such that any inherent side bias was cancelled out within groups. On day 2, mice were connected to a patch cable and placed into the unpaired chamber to begin the trial. 20 Hz, 5 ms red light stimulation (1 mW) was delivered whenever the center point of the mouse was in the paired chamber (controlled by Ethovision software, Noldus). The trial lasted for 20 min.

Intracranial self-stimulation: Mice were food restricted to 85% of ad libitum body weight to increase exploratory activity. Mice were connected to a patch cable and placed into an operant chamber (MedAssociates) for a 1-hour session each day for 4 days. Each session began with a 5 min “magazine” period in which both levers extended for 10 s, followed by lever retraction and delivery of a 3 s red laser light stimulation at 40 Hz (5 ms pulse duration, 5 mW). A press on the active lever during this 10 s period resulted in immediate lever retraction and red-light delivery. Mice received 10 magazine trials with a variable 30 s ITI. For the remainder of the session, levers extended and remained extended until a press was made on the active lever, which led to lever retraction and 3 s of 40 Hz light stimulation. Levers re-extended following an additional 2 s timeout period.

Following 4 days of training at 40 Hz, mice were tested once per day in a 30-min session with the stimulus frequency set to 5, 10, 20, or 40 Hz (one frequency per day, presented in a randomized order). Frequency testing sessions began with a 3-min magazine period with stimulation at the relevant frequency.

**Immunohistochemistry:**

Mice were perfused with 4% PFA. For confirmation of viral expression and fiber placements 50 µm frozen brain sections were collected and stained overnight. To quantify viral co-expression 30 µm frozen brain sections were collected and stained overnight. Antibodies used were Rabbit anti-HA (Sigma H6908, 1:1000), Chicken anti-GFP (AbCam 13970, 1:8000), and Mouse anti-DSRed (Clontech 632392, 1:2000). Secondary antibodies were from Jackson ImmunoResearch (1:250 dilution). Images were collected using a Keyence BZ-X710 fluorescent microscope and analyzed using ImageJ software.

**Statistics and Reproducibility:**

All data were analyzed for statistical significance using Prism software (GraphPad Prism 9). See Extended Data Table 2 for details on all statistical tests performed. The Geisser-Greenhouse correction was used to correct for unequal variability of differences in repeated-measures ANOVA tests. All behavioral assays were repeated in a minimum of two cohorts with similar replication of results. Littermates were randomly assigned to experimental groups and animals were tested in random order. Animals with missed viral injections or significant viral spread outside the targeted region were excluded from analyses.

**Supplementary Discussion**

Peptide and Fast Transmitter Co-Release

Many examples of neurotransmitter and neuropeptide co-expression have been identified in the CNS, with many potential configurations of convergent and divergent actions on downstream targets[3,4,29,30]. This includes LH neurons that release glutamate and orexin (both excitatory), which independently increase firing in the same downstream neurons on separate fast (glutamate) and slow (peptide) time scales[31]. Additionally, orexin/hypocretin neurons co-release stimulatory orexin and inhibitory dynorphin that can act either presynaptically or postsynaptically on different cells to facilitate inhibition or excitation[32], and can co-release GABA to directly inhibit postsynaptic cells[33], but how these actions coordinate circuit output is not clear. It has also been shown that orexin and dynorphin are co-released from the same vesicles to simultaneously inhibit or stimulate the same postsynaptic cell to antagonize each other's actions[34], but again how this functions operationally remains unresolved.

To our knowledge the mechanism we outline here of direct peptidergic excitation and di-synaptic fast transmitter disinhibition has not been previously described in a mammalian system. Intriguingly, this may prove to be a common circuit arrangement for inputs to the dopamine system, given the large number of GABAergic inputs to the VTA that synapse onto local GABA neurons[8], many of which release stimulatory peptides[35-37] that activate receptors enriched on dopamine neurons[28,38].

What are the potential advantages of this particular cooperative circuit organization? GABAergic disinhibition is a powerful mechanism for time-locked control of dopamine neuron firing[39], which is critical for regulating motivated behavior. Additionally, because we observed increasing peptidergic influence with increasing stimulation frequency, this circuit arrangement may serve to amplify bursts of activity that rise above steady state baseline firing[40]. The slower, longer-lasting calcium signal triggered by Ntsr1 activation in dopamine neurons may directly lead to a prolonged increase in firing and dopamine release during this period, and may also extend the window for coincidence detection of multiple inputs, facilitating plasticity and cue-reward association[41]. Notably, we found that the calcium signal in dopamine neurons following head entry lasted much longer (>20 s) than the calcium signal in LH-Nts neurons (~5 s), and the slow decay of this signal was significantly impacted by mutagenesis of *Ntsr1*. How this prolonged calcium signal translates into the specific dynamics of dopamine firing and downstream dopamine release remains to be determined.

Effects on calcium signals during behavior

Changes to the amplitude and kinetics of the dopamine neuron calcium signals were more subtle in behaving animals than following direct stimulation of LH-Nts neurons, likely due to the complexity of the activity pattern in LH-Nts neurons (ramp up and down as opposed to a square pulse) and to intact signaling from other inputs to the VTA that are almost certainly active during this behavior and are contributing to the calcium signal. We did not observe significant changes to the DA neuron calcium signals during the lever press or cue periods, which is consistent with these responses being driven primarily by other inputs to the VTA. This also likely explains why we did not observe impaired lever pressing during this task. This underscores the complexity of reward encoding by the dopamine system, which receives inputs from more than two dozen different brain regions.

We did observe that, in well-trained animals, mutagenesis of *Vgat* and *Ntsr1* increased the latency to retrieve the reward pellet towards the end of the session. This could indicate that these animals reach satiety earlier than controls. Notably, we found that over the course of several weeks these animals also lose body weight, and that silencing LH-Nts neurons with expression of TeTox has the same effect. Previous studies have found that infusion of Nts into the VTA or activation of LH-Nts neurons temporarily decreases food intake[42,43], while global knockout of *Ntsr1* tends to increase body weight[18,44]. While our data appear to contradict these findings, this is likely due to the circuit-specificity of our manipulation (as opposed to a global knockout), the onset of mutagenesis in adulthood (as opposed to in developing animals), and the sustained nature of our manipulation (as opposed to a brief infusion of Nts). A decrease in body weight is consistent with the decreased dopaminergic signaling during food reward that we observed, which may lead to reduced motivation for consumption. Further investigation of food intake and metabolism in these animals will shed light on the specific role of this circuit in long-term feeding behavior.

Potential for collateral effects

A potential caveat of our double disconnect strategy is disruption of signaling in collateral LH projections or from other VTA Nts inputs. While targeting *Ntsr1* in dopamine neurons will block signaling from all sources of Nts, this is unlikely to have a major impact on our stimulated photometry and optogenetic experiments, given that we are specifically activating only LH-Nts neurons. It is also possible that we are affecting GABA release in other target regions of LH-Nts neurons. However, our double disconnect strategy has an advantage in that both GABA and Nts signaling are simultaneously disrupted only at LH to VTA synapses; other collaterals still have one or the other element intact. Given that our optogenetic behavioral manipulations were stimulations of LH-Nts terminals within the VTA, and that disruption of Nts from LH neurons in had the same effect as disruption of Ntsr1, we find it most likely that the majority of effects we observed are mediated by direct LH to VTA connectivity. However, we cannot completely rule out a contribution from activation of indirect circuits.

Role of glutamate release

Although previous studies[6,7], as well as our own *in situ* hybridization data, indicate that the majority of LH-Nts neurons release GABA and only a small fraction release glutamate, co-release of GABA from these neurons has previously been ignored in favor of a glutamate-centric model[13], which seems more logical on its face due to the stimulatory actions of the neurotransmitter and neuropeptide. While we did detect a low level of glutamatergic connectivity from LH-Nts neurons to VTA neurons, we find it unlikely that glutamate release from LH-Nts neurons is a major driving force of dopamine neuron activation in this circuit. LH-Glu neurons as a whole synapse most strongly on non-dopamine VTA neurons[8,14], and stimulation of these neurons is behaviorally aversive and has been shown to decrease dopamine release[14] (but see [45] for evidence of direct LH glutamatergic activation of a specific subset of DA neurons). Though some residual calcium signal remained in our stimulated photometry experiment following mutagenesis of both *Vgat* and *Ntsr1*, given the circuit organization this is more likely due to incomplete mutagenesis than to activation by glutamate.

An additional technical consideration that argues against a significant role for glutamate in this LH-Nts to VTA circuit, which we noted previously[8], is the unequal distribution of functional LH to VTA glutamatergic connectivity along the anterior-posterior axis of the LH. The LH is a large structure that extends along the AP axis from -0.34 mm to -2.8 mm from bregma[46]. Previous findings of strong glutamatergic connectivity from the LH to the VTA have universally targeted the anterior portion of the LH (Nieh et al.: -0.4 to -0.8 mm, Kempadoo et al.: -0.4 mm, de Jong et al.: -0.8 mm, Soden et al. -0.6 mm). For this paper we targeted the central LH (-1.25 mm), which contains the largest population of Nts neurons. As we reported previously[8], we detected minimal glutamatergic LH to VTA connectivity using these coordinates, consistent with our findings specifically in LH-Nts neurons.

### Extended data figures and legends

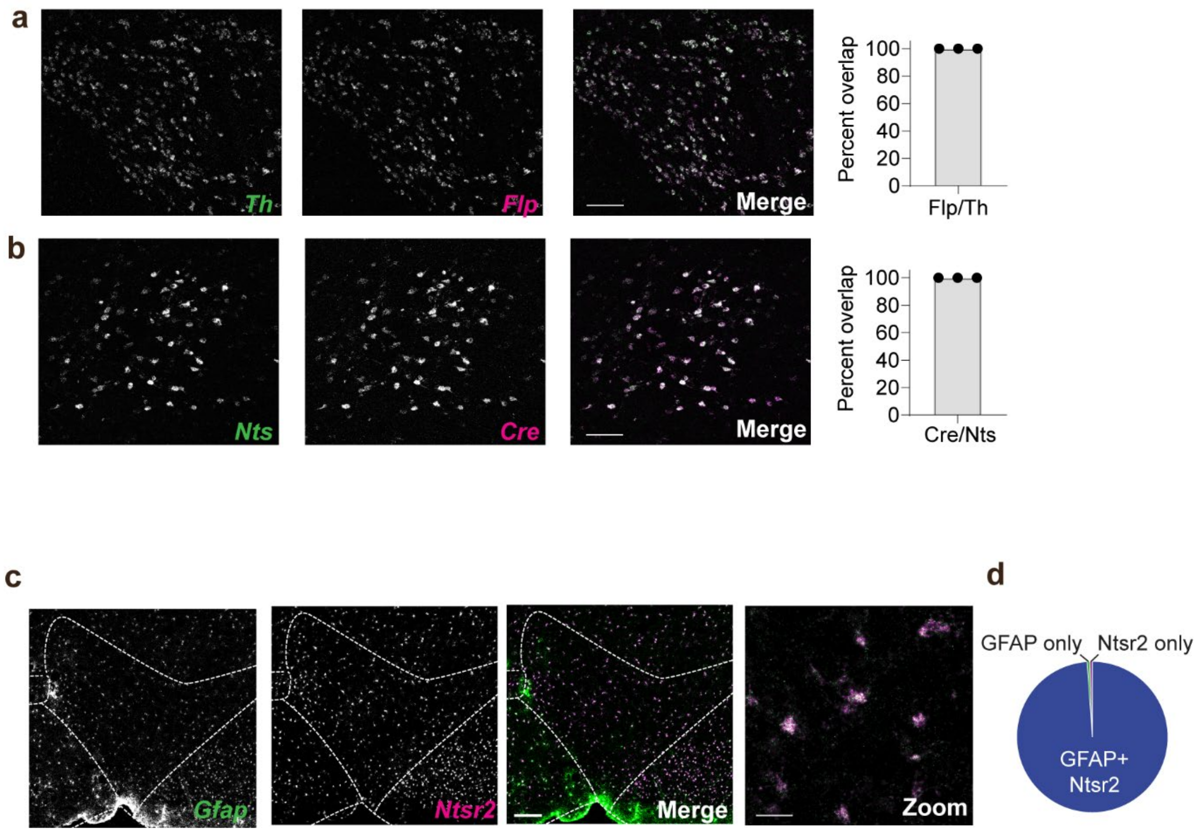


**Extended Data Fig. 1. Confirmation of specific recombinase expression and analysis of *Ntsr2* expression. (a)** Example images of *in situ* hybridization for *Th* and *Flp* recombinase in the VTA of Nts-Cre::Th-Flp mice, and percentage of *Flp*+ cells that expressed *Th* (n=9 sections from N=3 mice, 4332 total *Flp*+ cells, scale bar=100 µm). **(b)** Example images of *in situ* hybridization for *Nts* and *Cre* in the LH of Nts-Cre::Th-Flp mice, and percentage of *Cre*+ cells that expressed *Nts* (n=7 sections from N=3 mice, 1058 total *Cre*+ cells). **(c)** *In situ* hybridization for the astrocyte marker *Gfap* and *Ntsr2* in the VTA. Scale bars = 150 µm (greyscale images) and 25 µm (zoom). (**d**) Quantification of *Ntsr2* and *Gfap* overlap in the VTA (n=6 sections from N=2 mice). See Extended Data Table 1 for cell counts.

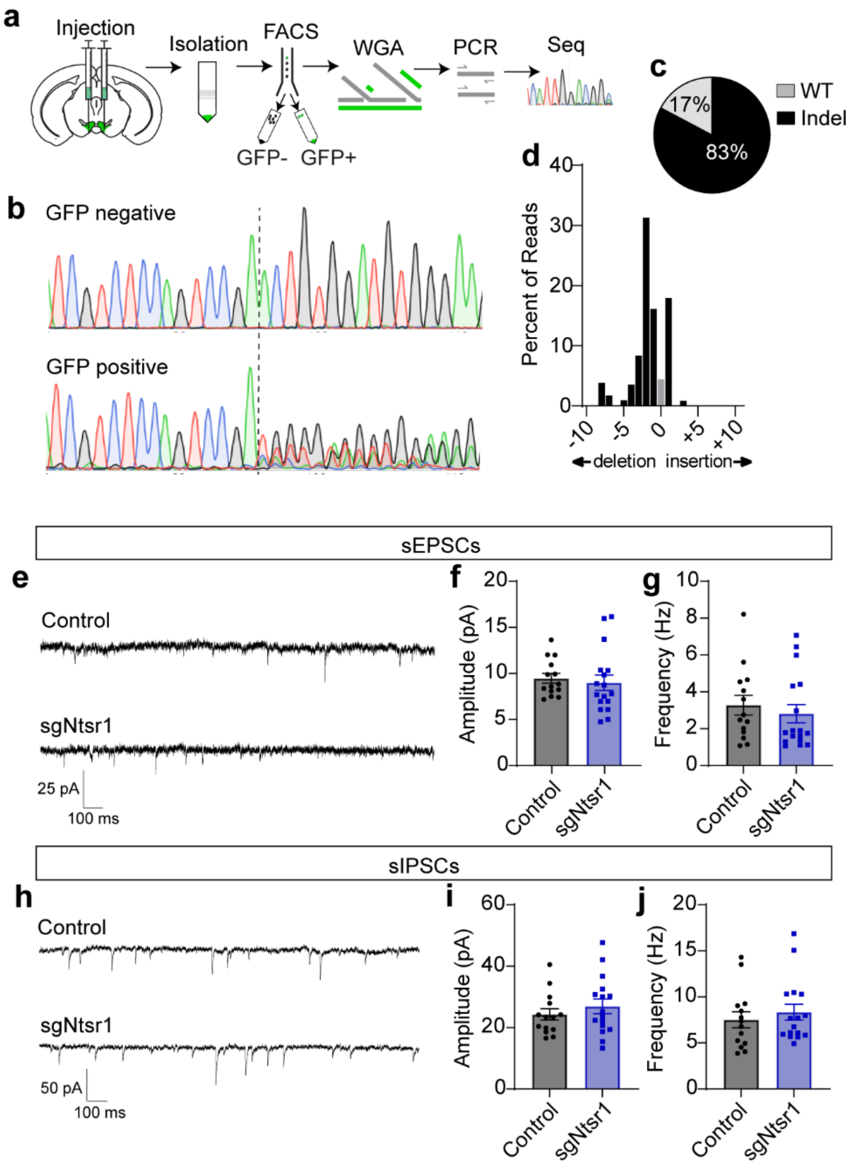


**Extended Data Fig. 2. Validation of AAV1-FLEXfrt-SaCas9-U6-sgNtsr1. (a)** Schematic of procedure for isolating GFP positive nuclei infected with sgNtsr1 CRISPR virus and nuclear envelope marker KASH-GFP followed by targeted sequencing of the intended cut site. **(b)** Sanger sequencing of pooled GFP negative and positive nuclei showing a disruption in the sequence at the intended cut site (dashed line) in GFP positive nuclei only. **(c)** Estimated percentage of WT and mutated (Indel) reads in GFP positive nuclei based on TIDE analysis. **(d)** Estimated percentage of each insertion and deletion mutation based on TIDE analysis. **(e)**

Example traces of spontaneous EPSCs recorded from VTA DA neurons expressing a control CRISPR or sgNtsr1. **(f-g)** sEPSC amplitude (**f**) and frequency (**g**) (control n=14 cells from N=2 mice, sgNtsr1 n=17 cells from N=2 mice). **(h)** Example traces of spontaneous IPSCs recorded from VTA DA neurons expressing a control CRISPR or sgNtsr1. **(f-g)** sIPSC amplitude (**f**) and frequency (**g**) (control n=14 cells from N=2 mice, sgNtsr1 n=16 cells from N=2 mice, scale bar=100 μm). Data are presented as mean ± S.E.M.

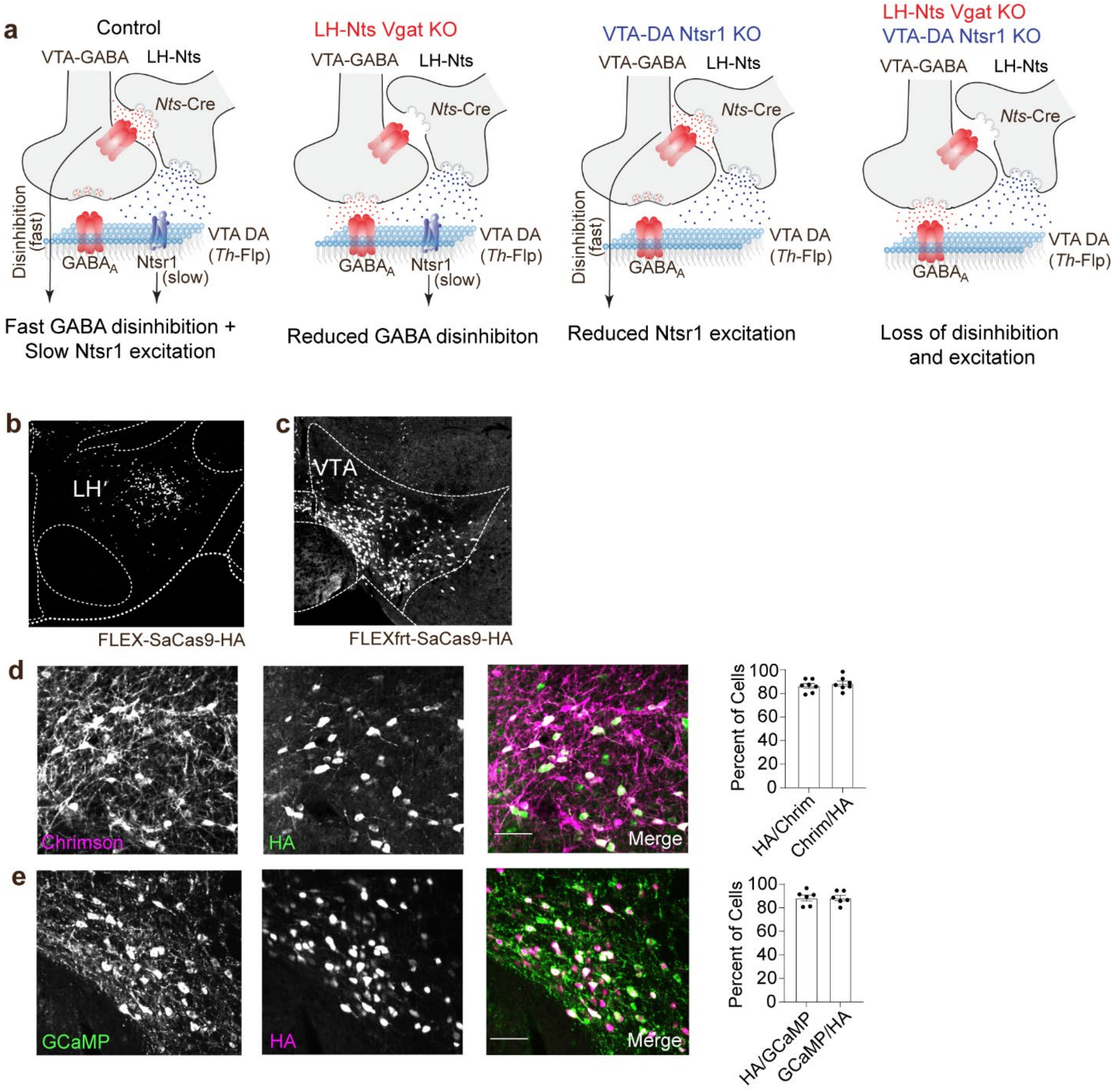


**Extended Data Fig. 3: Genetic strategy for anatomical disconnect and validation of viral co-expression.** (**a**) Cartoon illustrating hypothetical co-release of GABA and Nts from LH-Nts inputs to the VTA (Note: the axo-axonic connection is presented for simplicity and is not meant to reflect the only type of synapse involved in co-release). Inactivation of Vgat from LH-Nts neurons (LH-Nts GABA KO) is predicted to blunt disinhibition. Inactivation of Ntsr1 (VTA-DA Ntsr1 KO) is predicted to prevent Nts signaling onto VTA-DA neurons and blunt slow depolarization. Loss of both Vgat and Ntsr1 (LH-Nts GABA KO + VTA-DA Ntsr1 KO) is predicted to blunt both fast and slow depolarization of the VTA-DA neurons following activation of LH-Nts inputs. **(b)** Cre-dependent expression of SaCas9 in LH-Nts neurons from AAV1-FLEX-SaCas9-U6-sgVgat detected by immunostaining for HA tag. (**c**) Flp-dependent expression of SaCas9 in VTA-DA neurons from AAV1-FLEXfrt-SaCas9-U6-sgNtsr1 detected by immunostaining for HA tag. **(d)** Example images and quantification of co-expression of AAV1-FLEX-Chrimson-tdTomato and AAV1-FLEX-SaCas9-U6-sgVgat in the LH of Nts-Cre mice (n=7 sections from N=6 mice). **(e)** Example images and quantification of co-expression of AAV1-FLEXfrt-GCaMP6m and AAV1-FLEXfrt-SaCas9-U6-sgNtsr1 in the VTA of Th-Flp mice (n=6 sections from N=3 mice). Data are presented as mean ± S.E.M.

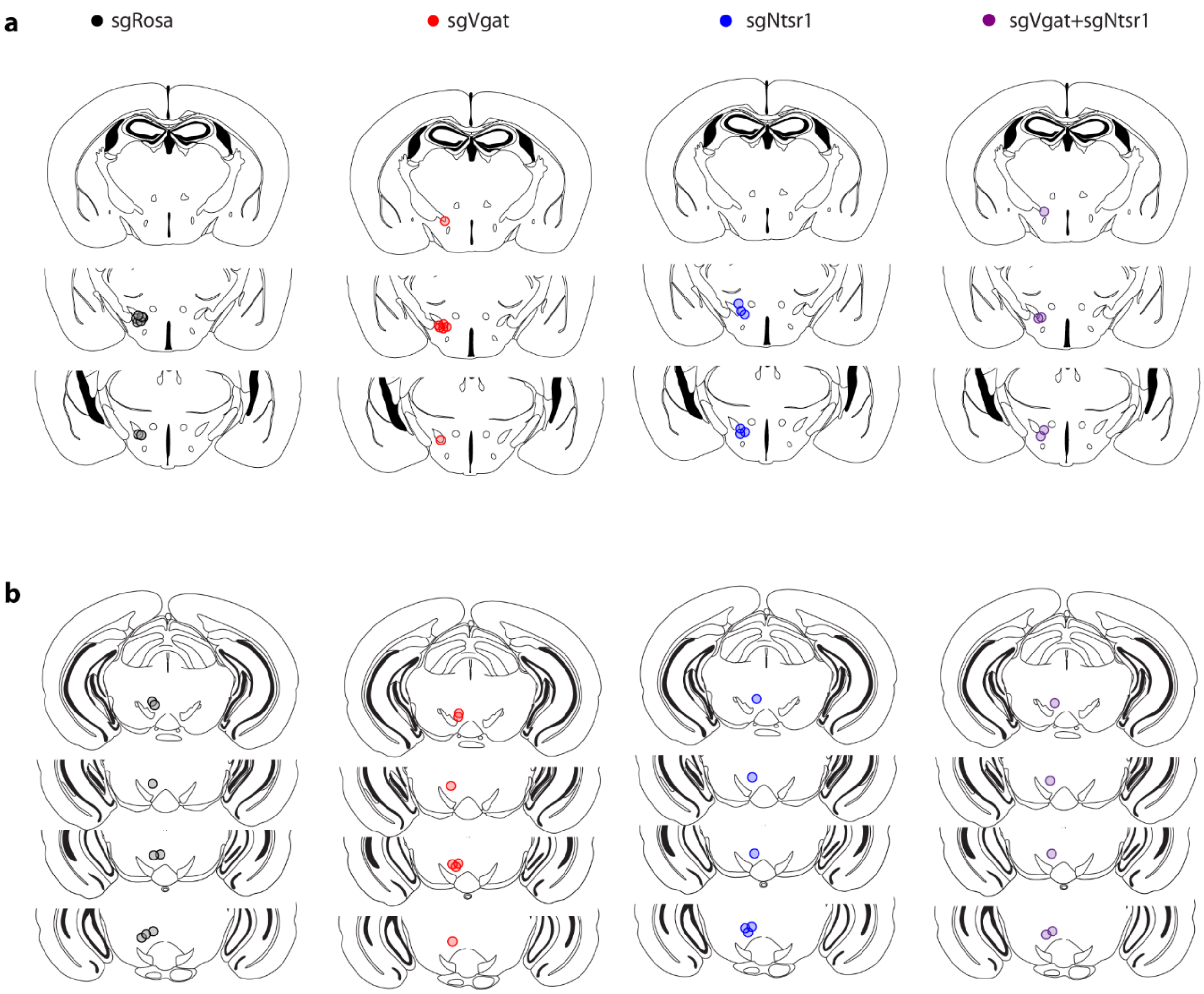


**Extended Data Fig. 4: Fiber placements for stimulation photometry experiments. (a)**

Placements for stimulatory fibers in LH. **(b)** Placements for recording fibers in VTA.

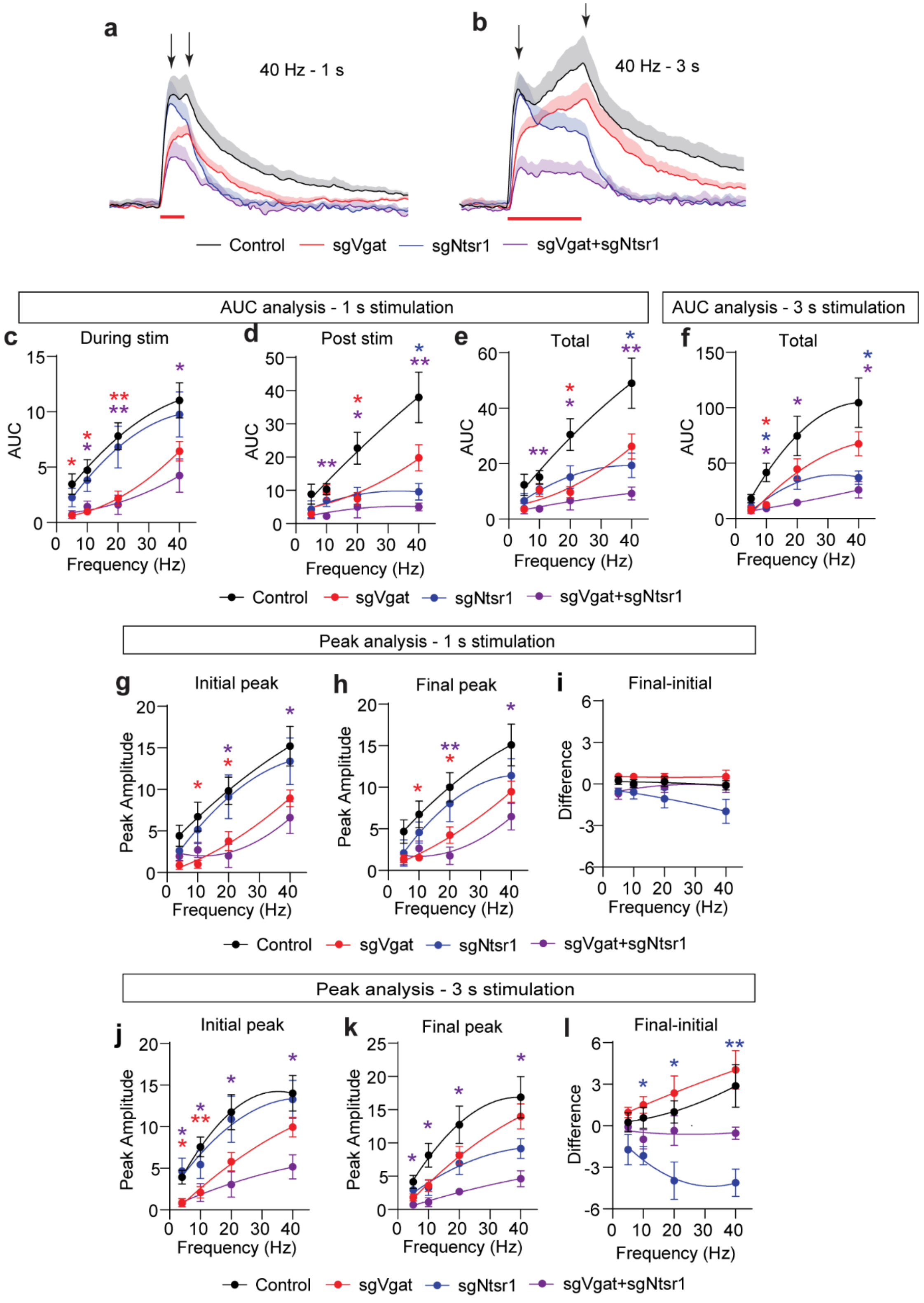


**Extended Data Fig. 5. Analysis of evoked GCaMP6m signals in VTA-DA neurons. (a-b)**

Average GCaMP6m fluorescence (Z-score) during and after 1 s (**a**) and 3 s (**b**) stimulation at 40

Hz. Arrows indicate initial and final peaks at the onset and offset of the stimulation, respectively. (**c**) Average area under the curve (AUC) of the Z-scored GCaMP6m fluorescence during 1 s stimulation at different frequencies. (**d**) Average AUC for the 10 s following 1 s stimulation at different frequencies. (**e**) Average AUC during and after 1 s stimulation at different frequencies. (**f**) Average AUC during and after 3 s stimulation at different frequencies. (**g**) Average initial peak amplitude (Z-score) of GCaMP6m fluorescence during 1 s stimulation at different frequencies. (**h**) Average final peak amplitude (Z-score) of GCaMP6m fluorescence during 1 s stimulation at different frequencies. (**i**) Difference between final and initial peak amplitude (final-initial) of GCaMP6m fluorescence during 1 s stimulation at different frequencies. (**j**) Average initial peak amplitude (Z-score) of GCaMP6m fluorescence during 3 s stimulation at different frequencies. (**k**) Average final peak amplitude (Z-score) of GCaMP6m fluorescence during 3 s stimulation at different frequencies. (**l**) Difference between final and initial peak amplitude (final-initial) of GCaMP6m fluorescence during 3 s stimulation at different frequencies. (Control N=8, sgVgat N=,7 sgNtsr1 N=6, sgVgat+sgNtsr1 N=5. *P<0.05; **P<0.01; ***P<0.001; red asterisk sgVgat vs. control; blue asterisk sgNts vs. control; purple asterisk sgVgat+sgNtsr1 vs. control.) Data are presented as mean ± S.E.M.

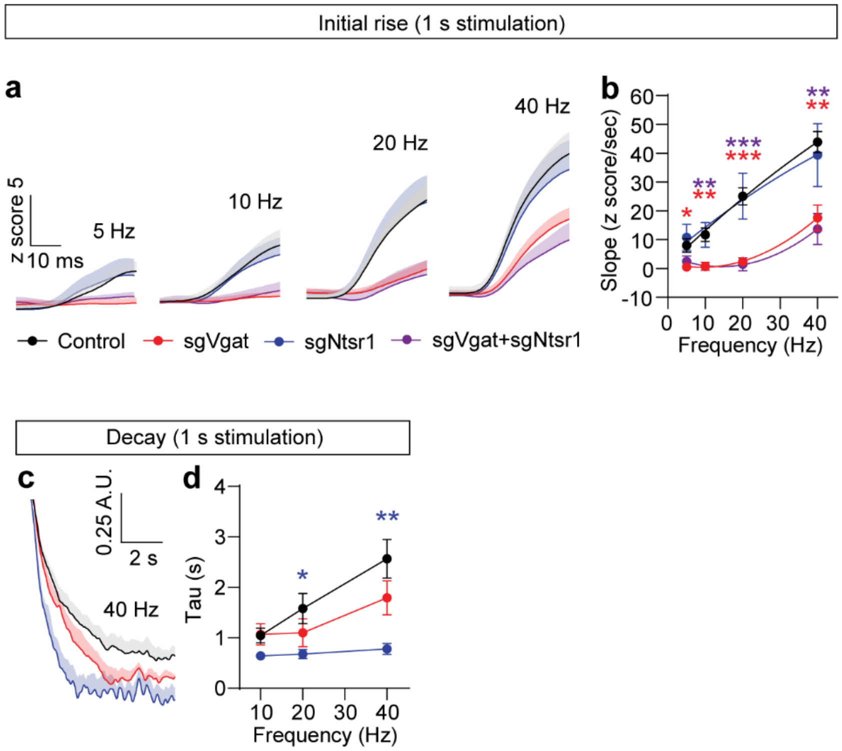


**Extended Data Figure 6. Analysis of evoked GCaMP6m signals in VTA-DA neurons. (a)** Average GCaMP6m fluorescence during the onset of 1 s optical stimulation. (**b**) Average slope of the initial rise in GCaMP6m fluorescence at different stimulus frequencies with 1 s optical stimulation. (**c**) Average GCaMP6m fluorescence decay following termination of 1 s stimulation, normalized to peak. (**d**) Average decay time constant of GCaMP6m fluorescence at different stimulus frequencies for 1 s. Note: Analysis of 5 Hz was omitted due to lack of signal in sgVgat mice, and sgVgat+sgNtsr1 mice were excluded from analysis due to the low amplitude of the evoked fluorescence. (Control N=8, sgVgat N=,7 sgNtsr1 N=6, sgVgat+sgNtsr1 N=5. *P<0.05; **P<0.01; ***P<0.001; red asterisk sgVgat vs. control; blue asterisk sgNts vs. control; purple asterisk sgVgat+sgNtsr1 vs. control). Data are presented as mean ± S.E.M.

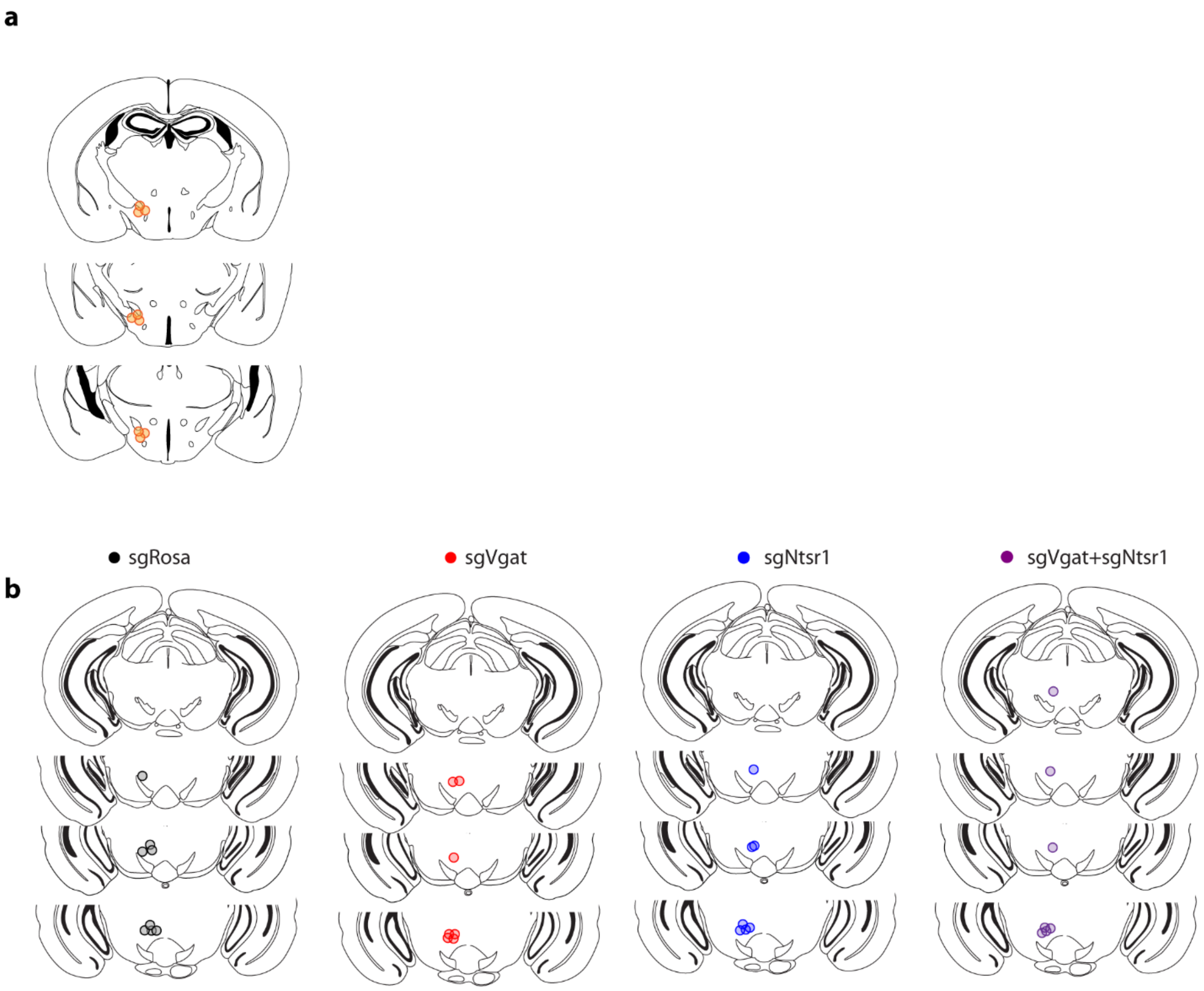


**Extended Data Figure 7. Fiber placements for behavioral photometry experiments. (a)**

Placements for recording fibers in LH. **(b)** Placements for recording fibers in VTA.

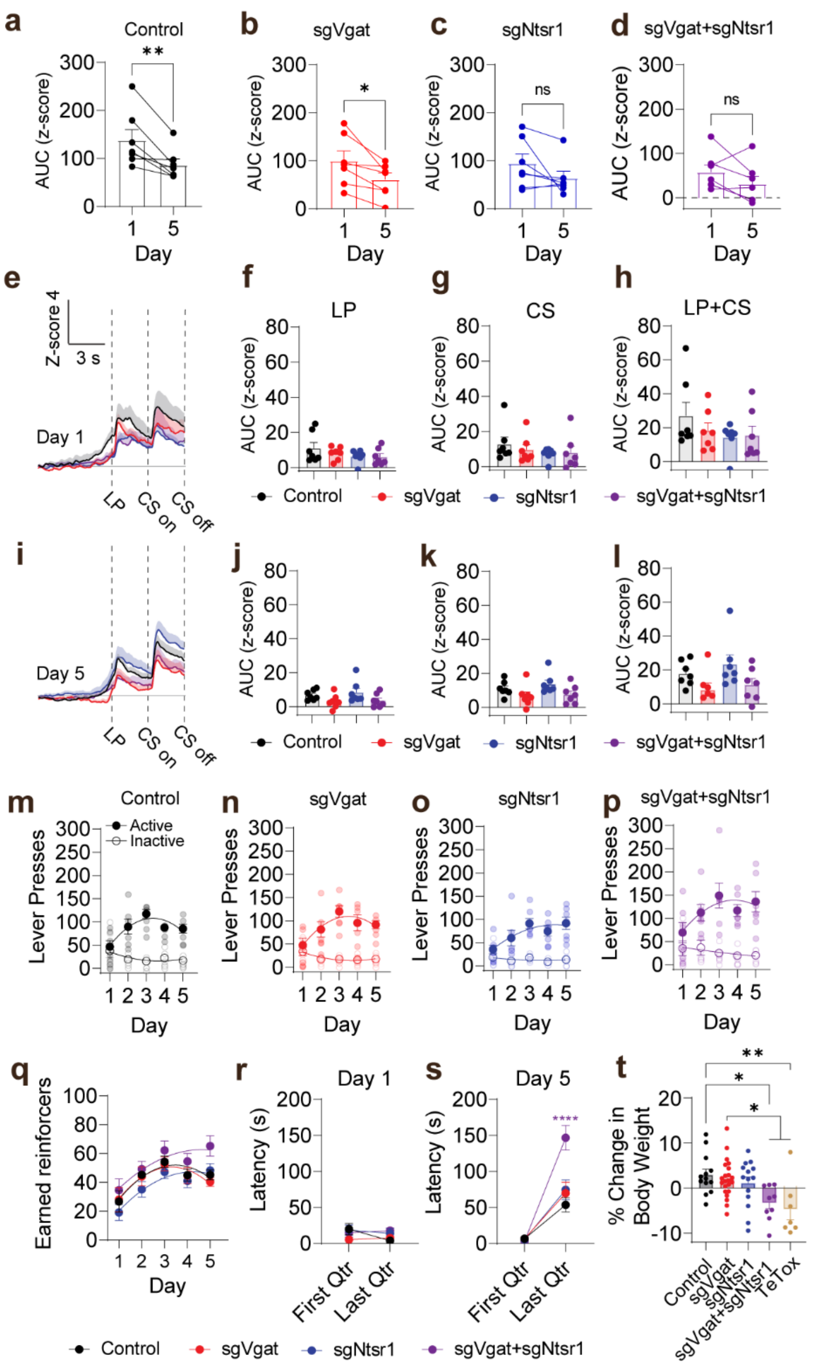


**Extended Data Figure 8. Additional analysis of behavioral photometry data. (a-d)** AUC of the z-scored fluorescence for the 20 s following head entry on days 1 and 5 in the indicated groups (N=7 mice per group, *p<0.05, **P<0.01). **(e)** Average GCaMP6m fluorescence on day 1 aligned to rewarded lever presses. **(f)** AUC during the 3 s following the lever press on day 1. **(g)** AUC during the CS period on day 1. **(h)** Summed AUC for the lever press and CS period on day

1. **(i)** Average GCaMP6m fluorescence on day 5 aligned to rewarded lever presses. **(j)** AUC during the 3 s following the lever press on day 5. **(k)** AUC during the CS period on day 5. **(l)** Summed AUC for the lever press and CS period on day 5. **(m-p)** Active and inactive lever presses across training days in the indicated groups. **(q)** Earned reinforcers across training days. **(r-s)** Latency to make a head entry into the food hopper following pellet delivery in the first or last quarter of the session on day 1 (r) or day 5 (s), averaged across all trials from all animals. **(t)** Percent change in body weight from pre-surgery weight to 5 weeks post viral injection (Control N=13 mice; sgVgat N=18 mice; sgNtsr1 N=14 mice; sgVgat+sgNtsr1 N=9 mice; TeTox N=7 mice). Data are presented as mean ± S.E.M.

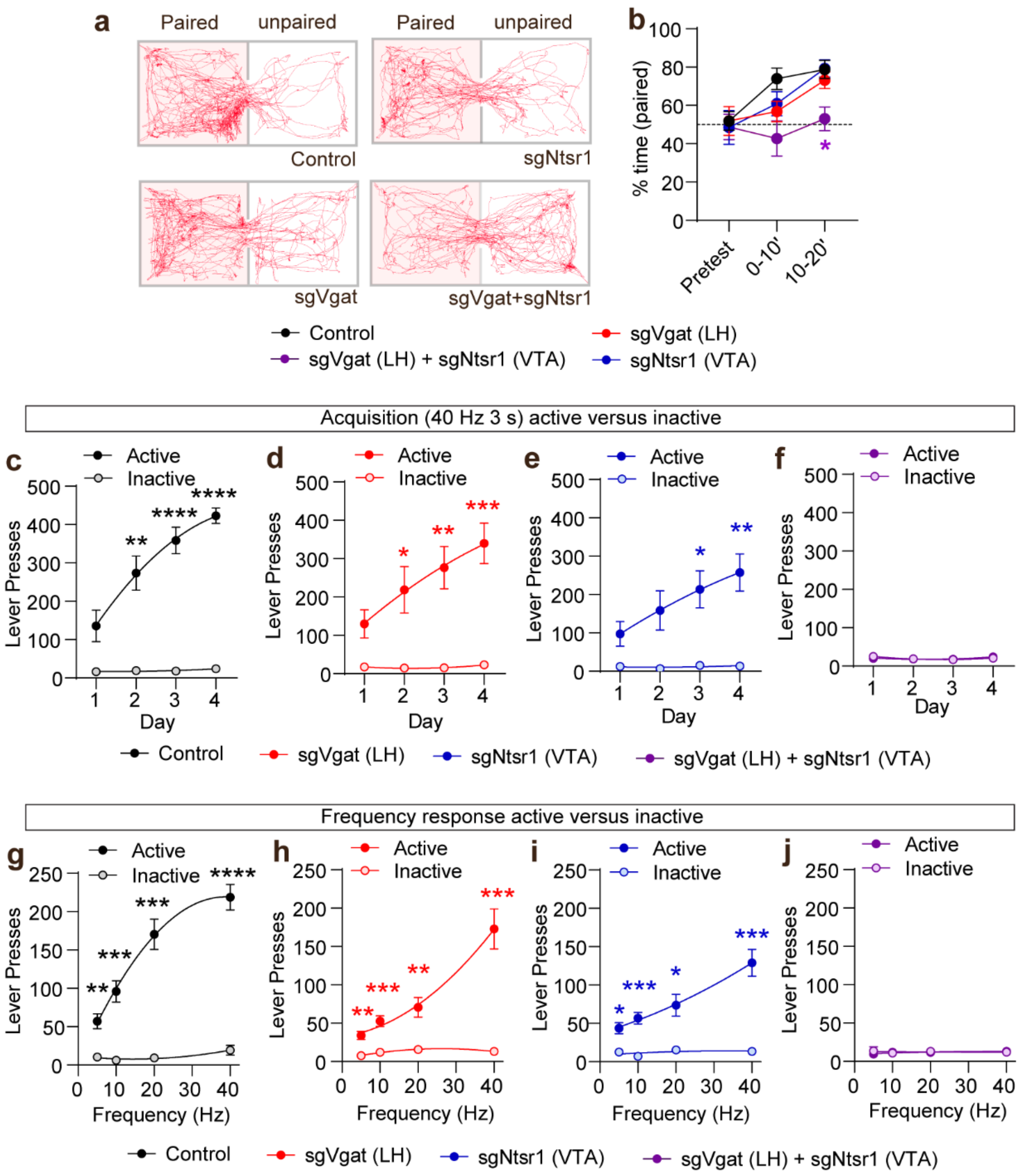


**Extended Data Figure 9. Coordinated actions of LH-Nts GABA release and VTA-DA Ntsr1 in behavioral reinforcement.** (**a**) Representative behavioral tracks during RTPP. (**b**) Average percent time in the light-paired chamber during 10 min pretest, and the first and last 10 min of RTPP (*P<0.05; control N=8, sgVgat N=,10 sgNtsr1 N=9, sgVgat+sgNtsr1 N=10; purple asterisk sgVgat+sgNtsr1 vs. control). (**c-f**) Average active and inactive lever presses for the four groups of mice during acquisition. (**g-j**) Average active and inactive lever presses for the four groups of mice during frequency response analysis (**c-j:** control N=9, sgVgat N=,10 sgNtsr1 N=8, sgVgat+sgNtsr1 N=10, *P<0.05; **P<0.01; ***P<0.001; ****P<0.0001). Data are presented as mean ± SEM.

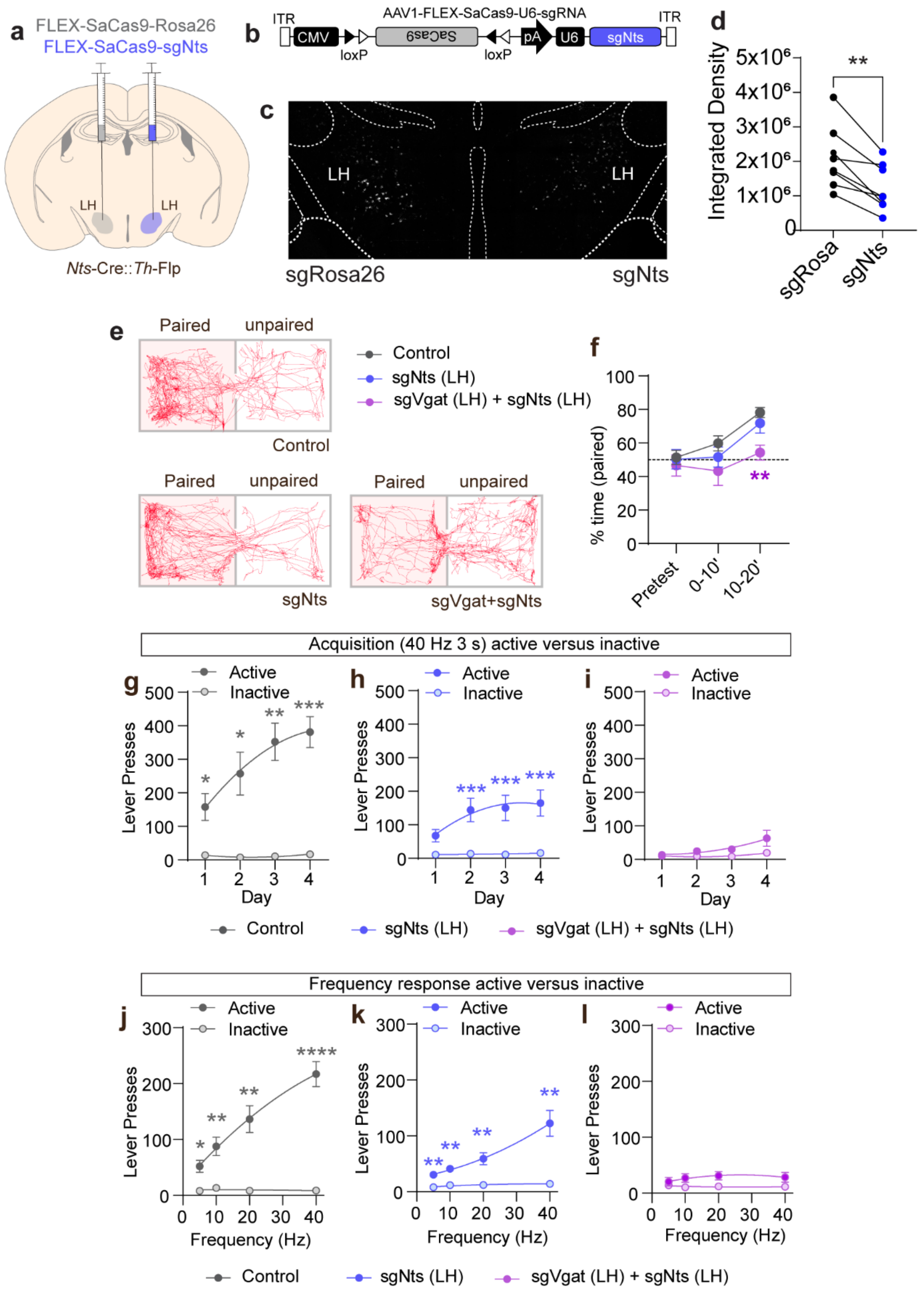

a
FLEX-SaCas9-Rosa26
FLEX-SaCas9-sgNts
LH
LH
Nts-Cre::Th-Flp
b
AAV1-FLEX-SaCas9-U6-sgRNA
ITR
CMV
SaCas9
pA
U6
sgNts
ITR
loxP
loxP
c
LH
LH
sgRosa26
sgNts
d
Integrated Density
**
sgRosa
sgNts
e
Paired
unpaired
Control
sgNts
sgVgat+sgNts
Control
sgNts (LH)
sgVgat (LH) + sgNts (LH)
f
% time (paired)
Pretest
0-10'
10-20'
Acquisition (40 Hz 3 s) active versus inactive
g
h
i
Active
Inactive
Lever Presses
Day
Control
sgNts (LH)
sgVgat (LH) + sgNts (LH)
Frequency response active versus inactive
j
k
l
Active
Inactive
Lever Presses
Frequency (Hz)
Control
sgNts (LH)
sgVgat (LH) + sgNts (LH)

**Extended Data Figure 10. Coordinated actions of LH-Nts GABA and Nts release in the VTA-DA for behavioral reinforcement.** (**a**) Schematic of unilateral injection of control virus (AAV1-FLEX-SaCas9-u6-sgRosa26 (grey)) and AAV1-FLEX-SaCas9-U6-sgNts (blue) into the LH of *Nts*-Cre mice. (**b**) Schematic of AAV1-FLEX-SaCas9-U6-sgNts viral vector. (**c**) Image of *in situ* hybridization for *Nts* in the sgRosa26 and sgNts injected sides of the LH. (**d**) Quantification of integrated pixel density of the fluorescent signal in the sgRosa26 and sgNts injected sides of the LH (**P<0.01; n=8 sections from N=4 mice). (**e**) Representative behavioral tracks during RTPP from the four groups of mice. (**f**) Average percent time in the light-paired chamber during 10 min pretest, and the first and last 10 min of RTPP (**P<0.01; purple asterisk sgVgat+sgNts vs. control; control N=12; sgNts=10; sgNts+sgVgat N=8). (**g-i**) Average active versus inactive lever presses for the three groups of mice during acquisition. (**j-l**) Average active versus inactive lever presses for the three groups of mice during frequency response analysis (**g-l** ****P<0.0001; ***P<0.001; **P<0.01; *P<0.05; control N=9; sgNts N=11; sgVgat+sgNts N=7). Data are presented as mean ± SEM.

**Extended Data Table 1: Cell counts.**

| Figure | N | Group | Cell Count | Percentage |
|---|---|---|---|---|
| 1b | n=9 sections from N=3 mice | **Inner Ring** | | |
| | | Vgat | 6742 | 65.9% |
| | | Vglut2 | 3352 | 32.7% |
| | | Vgat+Vglut2 | 139 | 1.4% |
| | | **Outer Ring** | | |
| | | Vgat+Nts | 2151 | 31.9% (of Vgat neurons) |
| | | Vglut2+Nts | 216 | 6.4% (of Vglut2 neurons) |
| | | Vgat+Vglut2+Nts | 48 | 34% (of Vgat+Vglut2 neurons) |
| 1c | n=9 sections from N=3 mice | Vgat+Nts | 2151 | 89.1% |
| | | Vglut2+Nts | 216 | 8.9% |
| | | Vgat+Vglut2+Nts | 48 | 2.0% |
| 1h | n=5 sections from N=2 mice | Th | 21 | 1.1% |
| | | Ntsr1 | 21 | 1.1% |
| | | Th+Ntsr1 | 1843 | 97.8% |
| Extended Fig. 1d | n=6 sections from N=2 mice | Gfap | 8 | 0.5% |
| | | Ntsr2 | 10 | 0.4% |
| | | Gfap+Ntsr2 | 1949 | 99.1% |

**Extended Data Table 2. Summary of statistical tests performed.**

| Figure | | Test | N | Statistics | P | Posttest | p (post) |
|---|---|---|---|---|---|---|---|
| **Main figure statistics** | | | | | | | |
| 2 | a | Unpaired t-test | $N_{control}$=11 cells; $N_{sgVgat}$=7 cells | | 0.0039 | | |
| | b | Unpaired t-test | $N_{control}$ =10 cells; $N_{sgNtsr1}$=8 cells | | 0.0173 | | |
| | f | Two-way RM ANOVA | $N_{control}$=8 mice; $N_{sgVgat}$=7 mice; $N_{sgNtsr1}$=6 mice; $N_{sgVgat+sgNtsr1}$=5 mice | Effect of frequency: $F_{(2.069, 45.52)}$=41.71, Effect of virus: $F_{(3, 22)}$ =6.790 | Effect of frequency: P<0.0001, Effect of virus: P=0.0021 | Dunnett's multiple comparisons test | *p<0.05, **p<0.01 (compared to control) |
| | g | Two-way RM ANOVA | $N_{control}$=8 mice; $N_{sgVgat}$=7 mice; $N_{sgNtsr1}$=6 mice; $N_{sgVgat+sgNtsr1}$=5 mice | Interaction $F_{(9,66)}$=3.061 | P=0.0040 | Dunnett's multiple comparisons test | *p<0.05 (compared to control) |
| | i | Two-way RM ANOVA | $N_{control}$=8 mice; $N_{sgVgat}$=7 mice; $N_{sgNtsr1}$=6 mice; $N_{sgVgat+sgNtsr1}$=5 mice | Interaction: $F_{(9,66)}$=2.404 | Interaction: P=0.0201 | Dunnett's multiple comparisons test | *p<0.05, **p<0.01, ***p<0.001 (compared to control) |

| | | | | | | | |
|---|---|---|---|---|---|---|---|
| | k | Two-way RM ANOVA | $N_{control}$=8 mice; $N_{sgVgat}$=7 mice; $N_{sgNtsr1}$=6 mice; | Effect of frequency: $F_{(1.410, 25.39)}$= 8.331, Effect of virus: $F_{(2, 18)}$ =5.817 | Effect of frequency: P=0.0039, Effect of virus: P=0.0113 | Dunnett's multiple comparisons test | *p<0.05 (compared to control) |
| 3 | d | One-sample t-test | N=9 mice | | Day 1 CS: P=0.0423; Day 1 HE P<0.0001; Day 5 HE P=0.0002 | | |
| | g | One-way ANOVA | N=7 mice/group | $F_{(3,24)}$=2.910 | P=0.0552 | Tukey's multiple comparisons test | *p<0.05 |
| | h | One-way ANOVA | N=7 mice/group | $F_{(3,24)}$=3.012 | P=0.0499 | Tukey's multiple comparisons test | *p<0.05 |
| | i | One-way ANOVA | N=7 mice/group | $F_{(3,24)}$=3.130 | P=0.0443 | Tukey's multiple comparisons test | *p<0.05 |
| | l | One-way ANOVA | N=7 mice/group | $F_{(3,24)}$=3.566 | P=0.0290 | Tukey's multiple comparisons test | *p<0.05 |
| | m | One-way ANOVA | N=7 mice/group | $F_{(3,24)}$=9.814 | P=0.0002 | Tukey's multiple comparisons test | *p<0.05, **p<0.01, ***p<0.001 |
| 4 | b | One-way ANOVA | $N_{control}$=8 mice; $N_{sgVgat}$=10 mice; $N_{sgNtsr1}$=9 mice; $N_{sgVgat+sgNtsr1}$=10 mice | $F_{(3,33)}$=6.150 | P=0.0019 | Tukey's multiple comparisons test | *p<0.05, **p<0.01 |
| | c | Two-way RM ANOVA | $N_{control}$=9 mice; $N_{sgVgat}$=10 mice; $N_{sgNtsr1}$=8 mice; $N_{sgVgat+sgNtsr1}$=10 mice | Interaction $F_{(9,99)}$=5.296 | P<0.0001 | Dunnett's multiple comparisons test | *p<0.05, **p<0.01, ***p<0.001 (compared to control) |
| | d | Two-way RM ANOVA | $N_{control}$=9 mice; $N_{sgVgat}$=10 mice; $N_{sgNtsr1}$=8 mice; $N_{sgVgat+sgNtsr1}$=10 mice | Interaction $F_{(9,99)}$=13.38 | P<0.0001 | Dunnett's multiple comparisons test | *p<0.05, **p<0.01, ***p<0.001 (compared to control) |
| | f | One-way ANOVA | $N_{control}$=12 mice; $N_{sgNts}$=10 mice; $N_{sgVgat+sgNts}$=8 mice | $F_{(2,27)}$=6.996 | P=0.0036 | Tukey's multiple comparisons test | *p<0.05, **p<0.01 |

| | | | | | | | |
|---|---|---|---|---|---|---|---|
| | g | Two-way RM ANOVA | $N_{control}$=9 mice; $N_{sgNts}$=11 mice; $N_{sgVgat+sgNts}$=7 mice | Interaction $F_{(6,72)}$=3.897 | P=0.002 | Dunnett's multiple comparisons test | *p<0.05, **p<0.01, ***p<0.001 (compared to control) |
| | h | Two-way RM ANOVA | $N_{control}$=9 mice; $N_{sgNts}$=11 mice; $N_{sgVgat+sgNts}$=7 mice | Interaction $F_{(6,72)}$=8.662 | P<0.0001 | Dunnett's multiple comparisons test | *p<0.05, **p<0.01, ***p<0.001 (compared to control) |
| **Extended Data Figure Statistics** | | | | | | | |
| E5 | c | Two-way RM ANOVA | $N_{control}$=8 mice; $N_{sgVgat}$=7 mice; $N_{sgNtsr1}$=6 mice; $N_{sgVgat+sgNtsr1}$=5 mice | Effect of frequency: $F_{(1.822, 40.09)}$=55.84, Effect of virus: $F_{(3, 22)}$ =6.005 | Effect of frequency: P<0.0001, Effect of virus: P=0.0038 | Dunnett's multiple comparisons test | *p<0.05, **p<0.01 (compared to control) |
| | d | Two-way RM ANOVA | $N_{control}$=8 mice; $N_{sgVgat}$=7 mice; $N_{sgNtsr1}$=6 mice; $N_{sgVgat+sgNtsr1}$=5 mice | Interaction $F_{(9,66)}$=5.167 | P<0.0001 | Dunnett's multiple comparisons test | *p<0.05, **p<0.01 (compared to control) |
| | e | Two-way RM ANOVA | $N_{control}$=8 mice; $N_{sgVgat}$=7 mice; $N_{sgNtsr1}$=6 mice; $N_{sgVgat+sgNtsr1}$=5 mice | Interaction $F_{(9,66)}$=4.554 | P=0.0001 | Dunnett's multiple comparisons test | *p<0.05, **p<0.01 (compared to control) |
| | f | Two-way RM ANOVA | $N_{control}$=8 mice; $N_{sgVgat}$=7 mice; $N_{sgNtsr1}$=6 mice; $N_{sgVgat+sgNtsr1}$=5 mice | Interaction $F_{(9,66)}$=2.535 | P=0.0146 | Dunnett's multiple comparisons test | *p<0.05 (compared to control) |
| | g | Two-way RM ANOVA | $N_{control}$=8 mice; $N_{sgVgat}$=7 mice; $N_{sgNtsr1}$=6 mice; $N_{sgVgat+sgNtsr1}$=5 mice | Effect of frequency: $F_{(2.157, 47.46)}$=54.43, Effect of virus: $F_{(3, 22)}$ =4.166 | Effect of frequency: P<0.001, Effect of virus: P=0.0177 | Dunnett's multiple comparisons test | *p<0.05 (compared to control) |
| | h | Two-way RM ANOVA | $N_{control}$=8 mice; $N_{sgVgat}$=7 mice; $N_{sgNtsr1}$=6 mice; $N_{sgVgat+sgNtsr1}$=5 mice | Effect of frequency: $F_{(1.893, 41.64)}$=55.18, Effect of virus: $F_{(3, 22)}$ =4.582 | Effect of frequency: P<0.001, Effect of virus: P=0.0122 | Dunnett's multiple comparisons test | *p<0.05, **p<0.01 (compared to control) |
| | i | Two-way RM ANOVA | $N_{control}$=8 mice; $N_{sgVgat}$=7 mice; $N_{sgNtsr1}$=6 mice; $N_{sgVgat+sgNtsr1}$=5 mice | Effect of virus: $F_{(3, 22)}$ =4.094 | Effect of virus: P=0.0188 | Dunnett's multiple comparisons test | No significant post-hoc comparisons |
| | j | Two-way RM ANOVA | $N_{control}$=8 mice; $N_{sgVgat}$=7 mice; $N_{sgNtsr1}$=6 mice; $N_{sgVgat+sgNtsr1}$=5 mice | Effect of frequency: $F_{(2.163, 47.59)}$=34.94, Effect of virus: $F_{(3, 22)}$ =7.225 | Effect of frequency: P<0.001, Effect of virus: P=0.0015 | Dunnett's multiple comparisons test | *p<0.05, **p<0.01 (compared to control) |

| | | | | | | | |
|---|---|---|---|---|---|---|---|
| | k | Two-way RM ANOVA | $N_{control}$=8 mice; $N_{sgVgat}$=7 mice; $N_{sgNtsr1}$=6 mice; $N_{sgVgat+sgNtsr1}$=5 mice | Interaction: $F_{(9,66)}$=2.725 | P=0.0091 | Dunnett's multiple comparisons test | *p<0.05 (compared to control) |
| | l | Two-way RM ANOVA | $N_{control}$=8 mice; $N_{sgVgat}$=7 mice; $N_{sgNtsr1}$=6 mice; $N_{sgVgat+sgNtsr1}$=5 mice | Interaction: $F_{(9,66)}$=2.574 | P=0.0133 | Dunnett's multiple comparisons test | *p<0.05, **p<0.01 (compared to control) |
| E6 | b | Two-way RM ANOVA | $N_{control}$=8 mice; $N_{sgVgat}$=7 mice; $N_{sgNtsr1}$=6 mice; $N_{sgVgat+sgNtsr1}$=5 mice | Interaction $F_{(9,66)}$=3.485 | P=0.0014 | Dunnett's multiple comparisons test | *p<0.05, **p<0.01, ***p<0.001 (compared to control) |
| | d | Mixed-effects analysis | $N_{control}$=8 mice; $N_{sgVgat}$=7 mice*; $N_{sgNtsr1}$=6 mice*; | Effect of frequency: $F_{(1.627, 26.04)}$= 8.516, Effect of virus: $F_{(2, 18)}$ =7.760 | Effect of frequency: P=0.0025, Effect of virus: P=0.0037 | Dunnett's multiple comparisons test | *p<0.05, **p<0.01 (compared to control) |
| | *For panel 3d, mice with a peak z score <1 for a given stimulation frequency were excluded from analysis at that frequency, necessitating a mixed-effects analysis as opposed to a Two-way ANOVA. Mice excluded were: 3 sgVgat and 1 sgNtsr1 at 10 Hz | | | | | | |
| E8 | i | Paired t-test | N=7 mice/group | | P=0.0097 | | |
| | j | Paired t-test | N=7 mice/group | | P=0.0200 | | |
| | s | Two-way ANOVA | $N_{control}$=64 first qtr, 69 last qtr events; $N_{sgVgat}$=77 first qtr, 53 last qtr events; $N_{sgNtsr1}$=73 first qtr, 82 last qtr events; $N_{sgVgat+sgNtsr1}$= 113 first qtr, 78 last qtr events | Interaction: $F_{(3,601)}$=11.15 | Interaction: P<0.0001 | Dunnett's multiple comparisons test | ****p<0.0001 compared to control |
| | t | One-way ANOVA | $N_{control}$=13 mice; $N_{sgVgat}$=18 mice; $N_{sgNtsr1}$=14 mice; $N_{sgVgat+sgNtsr1}$=9 mice $N_{TeTox}$=7 mice | $F_{(4,60)}$=5.154 | P=0.0012 | Tukey's multiple comparisons | *p<0.05, **p<0.01 |
| E9 | b | Two-way RM ANOVA | $N_{control}$=8 mice; $N_{sgVgat}$=10 mice; $N_{sgNtsr1}$=9 mice; $N_{sgVgat+sgNtsr1}$=10 mice | Effect of time: $F_{(1.599, 52.76)}$= 15.72, Effect of virus: $F_{(3, 33)}$ =3.084 | Effect of time: P<0.0001, Effect of virus: P=0.0406, | Dunnett's multiple comparisons test | *p<0.05 (compared to control) |

| | | | | | | | |
|---|---|---|---|---|---|---|---|
| | c | Two-way RM ANOVA | N=9 mice | Interaction: $F_{(3,48)}$=15.98 | Interaction: P<0.0001 | Sidak's multiple comparisons test | **p<0.01, ****p<0.0001 |
| | d | Two-way RM ANOVA | N=10 mice | Interaction: $F_{(3,54)}$=11.35 | Interaction: P<0.0001 | Sidak's multiple comparisons test | *p<0.05, **p<0.01, ***p<0.001 |
| | e | Two-way RM ANOVA | N=8 mice | Interaction: $F_{(3,42)}$=6.752 | Interaction: P=0.0008 | Sidak's multiple comparisons test | *p<0.05, **p<0.01 |
| | g | Two-way RM ANOVA | N=9 mice | Interaction: $F_{(3,48)}$=35.47 | Interaction: P<0.0001 | Sidak's multiple comparisons test | **p<0.01, ***p<0.001, ****p<0.0001 |
| | h | Two-way RM ANOVA | N=10 mice | Interaction: $F_{(3,54)}$=22.48 | Interaction: P<0.0001 | Sidak's multiple comparisons test | **p<0.01, ***p<0.001 |
| | i | Two-way RM ANOVA | N=8 mice | Interaction: $F_{(3,42)}$=17.63 | Interaction: P<0.0001 | Sidak's multiple comparisons test | *p<0.05, ***p<0.001 |
| E 10 | d | Paired t-test | n=8 sections from N=4 mice | | P=0.0016 | | |
| | f | Two-way RM ANOVA | $N_{control}$=12 mice; $N_{sgNts}$=10 mice; $N_{sgVgat+sgNts}$=8 mice | Effect of time: $F_{(1.726, 46.61)}$= 12.61, Effect of virus: $F_{(2, 27)}$ =4.452 | Effect of time: P<0.0001, Effect of virus: P=0.0213, | Dunnett's multiple comparisons test | **p<0.01 (compared to control) |
| | g | Two-way RM ANOVA | N=9 mice | Interaction: $F_{(3,48)}$=11.84 | Interaction: P<0.0001 | Sidak's multiple comparisons test | *p<0.05, **p<0.01, ***p<0.001 |
| | h | Two-way RM ANOVA | N=11 mice | Interaction: $F_{(3,60)}$=4.918 | Interaction: P=0.0040 | Sidak's multiple comparisons test | ***p<0.001 |
| | i | Two-way RM ANOVA | N=7 mice | Effect of frequency: $F_{(1.143,13.72)}$=4.862, | Effect of frequency: P=0.0411 | Sidak's multiple comparisons test | No significant post-hoc comparisons |
| | j | Two-way RM ANOVA | N=9 mice | Interaction: $F_{(3,48)}$=31.70 | Interaction: P<0.0001 | Sidak's multiple comparisons test | *p<0.05, **p<0.01, ****p<0.0001 |

| | | | | | | | |
|---|---|---|---|---|---|---|---|
| | k | Two-way RM ANOVA | N=11 mice | Interaction: $F_{(3,60)}=11.35$ | Interaction: P<0.0001 | Sidak's multiple comparisons test | **p<0.01 |
